# iSBEM: An Open-Source Workflow for Automated ROI Targeting in Volume Electron Microscopy

**DOI:** 10.64898/2026.06.05.730298

**Authors:** Paolo Ronchi, Graham Ross, Alana Burrell, Joost de Folter, Yanneck Klenz, Nedal Darif, Fiona Young, Matthew Lawson, Jonas Albers, Tobias Pietz, Friedrich Frischknecht, Elizabeth Duke, Candice Roufosse, Lucy Collinson, Amy Strange, Yannick Schwab

**Author notes:** Equal contributions (Paolo Ronchi, Graham Ross, Alana Burrell).

## Abstract

Serial Block Face – Scanning Electron Microscopy (SBF-SEM) is a volume EM method suited to investigate the 3D architecture of tissues and even entire organisms at high resolution. However, imaging large volumes in their entirety is time-consuming and not always necessary. Many research projects have a focused interest in well-defined sub-regions of the samples. The targeting and acquisition of such regions of interest (ROIs) are however currently conducted in a manual way and require heavy involvement of experienced operators. We present a workflow and an original open-source software tool (iSBEM), which allow automated targeting of ROIs in a large tissue sample, based on X-ray microscopy (XRM) maps. After an initial ROI identification and registration of the XRM map with the sample mounted on the SBF-SEM stage, iSBEM takes over the control of the microscope, triggering high resolution acquisitions at defined ROI positions, with minimal user intervention. We demonstrate the approach on two biologically distinct specimens — malarial oocysts in infected mosquito midgut tissue, and immune cells in human kidney biopsies — achieving significant improvement in acquisition throughput relative to manual operations, without compromising targeting precision. We also showcase the workflow in a correlative light-Xray-electron microscopy setup, which allowed us to further improve the correct target definition.

## Introduction

The advent of volume electron microscopy (vEM) techniques over the past decade has revolutionized the ultrastructural analysis of biological specimens in three-dimensional space (Peddie and Collinson, 2014; Peddie et al., 2022; Titze and Genoud, 2016). Among these approaches, serial block-face scanning electron microscopy (SBF-SEM) has emerged as a key technique enabling the acquisition of large biological volumes at nanometer resolution. The original concept of an in-chamber miniaturized microtome was introduced by Leighton in 1981 and was later automated and further developed into a functional SBF-SEM platform by Denk and Horstmann in 2004 (Denk and Horstmann, 2004). During image acquisition, an integrated ultramicrotome repeatedly removes thin sections (typically 20–50 nm) from the resin block surface, after which the newly exposed block face is scanned by SEM to generate an electron micrograph. Iterative repetition of this cut-and-image cycle results in a three-dimensional ultrastructural dataset.

This large-volume acquisition strategy was initially established in neuroscience, where it was one of the approaches enabling the reconstruction of complex neural circuits and the generation of high-resolution connectomics datasets (e.g. Briggman and Bock, 2012; Wanner et al., 2016; Schmidt et al., 2017). However, SBF-SEM was rapidly adopted across the broader vEM community (Peddie et al., 2022) and has since been applied to a wide range of biological specimens, spanning from cultured cell mono-layers (Russell et al., 2017), single-celled parasites (Darif et al., 2026) to entire small organisms (Vergara et al., 2021), plants (Kittelmann et al., 2016), and human tissues (Randles et al., 2016). SBF-SEM is widely used because it combines high-resolution 3D ultrastructural imaging with the capacity to acquire large volumes that remain challenging for most other vEM techniques.

While vEM is often used for large-scale imaging, an alternative and increasingly relevant approach is region-of-interest (ROI) targeting. Instead of acquiring entire large volumes at high resolution, targeting workflows focus on identifying and imaging one or more small ROIs within a much larger sample volume. This is particularly important for complex biological samples where the structure of interest might only represent a small fraction of the specimen, and where blind acquisition would be inefficient, time-consuming, or not feasible (e.g. Karreman et al., 2016; Bishop et al., 2011; Maco et al., 2013; Kremer et al., 2021). In the particular case of SBF-SEM, sample preparation imposes specific limitations for targeting strategies. While post-embedding 3D correlative light and EM approaches (CLEM) have proven useful for FIB-SEM targeting (Ronchi et al., 2021), classical SBF-SEM heavy metal staining and resin embedding protocols do not allow fluorescence imaging of the final resin block. This prevents direct post-embedding fluorescence-based localization and requires alternative strategies for accurate navigation inside opaque resin blocks.

One of the most promising technologies to overcome this limitation is X-ray microscopy (XRM). XRM-based targeting approaches have been developed early on for transmission electron microscopy (TEM) workflows and block-face imaging applications. X-ray micro-computed tomography was used to visualize internal sample architecture, allowing sample quality validation and guiding subsequent sectioning or electron microscopy acquisition (Sengle et al., 2013; Bushong et al., 2015; Zheng et al., 2018; Hoffman et al., 2020; Handschuh et al., 2013). Importantly, XRM provides the ability to contextualize small vEM acquisitions within the surrounding sample volume, enabling high correlation specificity and accurate spatial navigation within resin-embedded samples. This capability has made XRM an increasingly attractive method for guiding vEM acquisition, particularly for workflows aiming to capture small volumes with high precision. Over the last years, multiple labs have implemented XRM-guided targeting pipelines for SBF-SEM and related vEM approaches (Laundon et al., 2023; Bosch et al., 2022; Burrell et al., 2025), and recent developments have introduced semi-automated strategies to streamline this process and reduce manual targeting effort (Meechan et al., 2022). Beyond its role as a targeting modality, XRM is also highly compatible with correlative imaging pipelines combining pre-embedding fluorescence microscopy and electron microscopy. This enables ROI selection based not only on ultrastructural context, but additionally on molecularly specific fluorescence information, thereby linking functional labelling to high-resolution vEM readouts (Karreman et al., 2016; Zhang et al., 2022; Svara et al., 2022).

Despite these advances, XRM-guided targeting remains technically demanding and is still not a routine workflow. Current approaches often require extensive manual handling steps, including sample remounting, repeated block trimming, and iterative re-imaging to confirm the ROI position (Laundon et al., 2023; Zhang et al., 2022). As a consequence, high-precision targeting remains time-intensive and can become a major bottleneck for projects aiming to systematically acquire multiple ROIs across large specimen volumes.

Here, we present “intelligent SBF-SEM” (iSBEM), an open-source napari (Sofroniew et al., 2022) plugin and integrated workflow designed to automate XRM-guided SBF-SEM acquisition of multiple ROIs within a single sample with minimal user intervention. We systematically compare different acquisition schemes, ranging from fully automated to supervised targeting strategies, and quantify the targeting accuracy achieved across multiple experimental conditions. By reducing manual handling and enabling reproducible multi-ROI acquisition, iSBEM substantially increases imaging throughput while maintaining high targeting precision. As a proof of concept, we successfully targeted malarial oocysts in infected mosquito midguts as well as immune cells in human kidney biopsies, demonstrating a highly robust workflow with a high accuracy rate in different sample types.

## Results

### Overview of the iSBEM workflow

The iSBEM workflow was developed to automate high-resolution imaging of defined subvolumes within large samples using SBF-SEM. The complete pipeline is illustrated in Figure 1 and detailed below. It leverages on a correlative X-ray and volume electron microscopy approach, where X-ray microscopy (XRM) is utilized to map samples, identify regions of interest (ROIs) based on morphological and anatomical features and predict their positions within resin embedded tissue (Fig. 1A). After mapping and ROI segmentation, samples are mounted in the SBF-SEM, where the cutting angle is fixed and the sample alignment cannot be adjusted. As the fixed orientation of the specimen may not match perfectly with the orientation of the XRM dataset, we designed a napari plugin (iSBEM) that first allows to register the XRM volume (together with the segmented ROIs) to the SBF-SEM coordinate system (Fig. 1A). The plugin then sends instructions to the microscope via its imaging software (the open-source SBEMimage (Titze et al., 2018)) to automate ROI acquisitions at their predicted locations (Fig. 1A,B). Figure 1B details the interplay between iSBEM and SBEMimage to drive the SBF-SEM targeted acquisitions: iSBEM allows the registration of XRM and EM Overviews (OVs), therefore defining ROI positions in the SBF-SEM coordinate system. These positions are then used by SBEMimage to create grids with tiles for imaging. Each grid can have different acquisition parameters (pixel size, tile size, etc), as defined in SBEMimage. During the acquisition, after each cut, iSBEM checks the predicted virtual slice for the presence of ROIs, and sends a command to SBEMimage for activating tiles at the corresponding position. The iSBEM workflow can be used for fully automated acquisitions (Fig. 1C, bottom arrow). Alternatively, the progression and accuracy of the registration can be monitored during the run, and eventually corrected, when necessary (Fig. 1C top).

**Figure 1.**
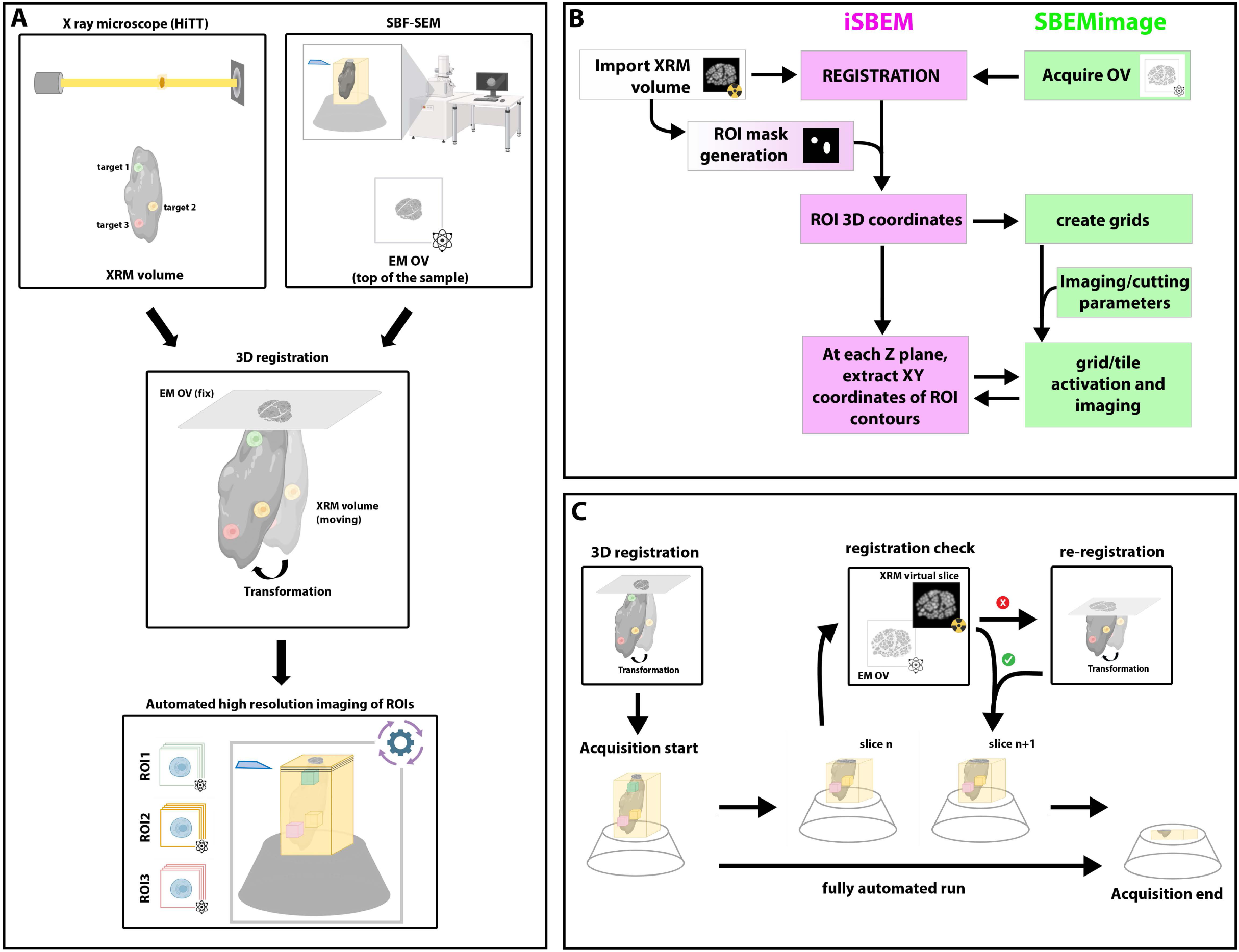
Strategy and iSBEM workflow. A. Automated SBF-SEM acquisition strategy: a tissue map is acquired by X-ray microscopy; targets are identified based on their morphological appearance in the XRM volume. The sample is then mounted in the SBF-SEM, where it is sectioned until the top of the tissue is exposed and a low resolution overview (OV) image is acquired. The XRM volume is then transformed to fit the sample orientation in the SBF-SEM. This registration defines the transformed position of the targets. Based on the ROI new coordinates, a command to acquire high resolution volumes in the right location is sent to the SBF-SEM acquisition software; the ROIs are acquired automatically with minimal non-relevant volumes around them. B. Interface between iSBEM and SBEMimage for automated ROI acquisition. Green rectangles highlight operations that are run by SBEMimage; Pink rectangles show operations done in iSBEM. White rectangles represent operations done with different software. ROI masks can be either generated within iSBEM or imported from a different software. C. iSBEM run supervision. After the initial registration, the acquisition can be run fully automated (lower arrow). Alternatively, registration accuracy can be checked at any point (slice n) during the run, comparing the current SBF-SEM OV to the corresponding XRM virtual slice. If necessary, the registration can be modified and the run can continue (slice n+1). Figure created using BioRender.com

### ROI definition

For the current study, we mapped our samples using X-ray High Throughput Tomography (HiTT) at the PETRA III synchrotron in Hamburg (Albers et al., 2024). Although the workflow can utilize any XRM data, HiTT provided superior throughput than lab source XRMs for identifying morphological features within the resin embedded tissue. After XRM mapping, the first step in the workflow is ROI definition. ROIs can be segmented offline with any available software and be imported as a mask stack in iSBEM. The plugin can also use integrated napari segmentation functionalities (see Materials and Methods). In the case studies presented here, we manually segmented ROIs using IMOD (Kremer et al., 1996) or Amira (Thermo Fisher Scientific) and imported the segmentation as binary TIFF masks in iSBEM. Regardless of the segmentation approach, masks can be further modified after import. For instance, ROIs can be dilated to provide buffer space in case of lower registration accuracy, or merged to prevent redundant scanning of closely spaced regions (Figure 2A).

**Figure 2.**
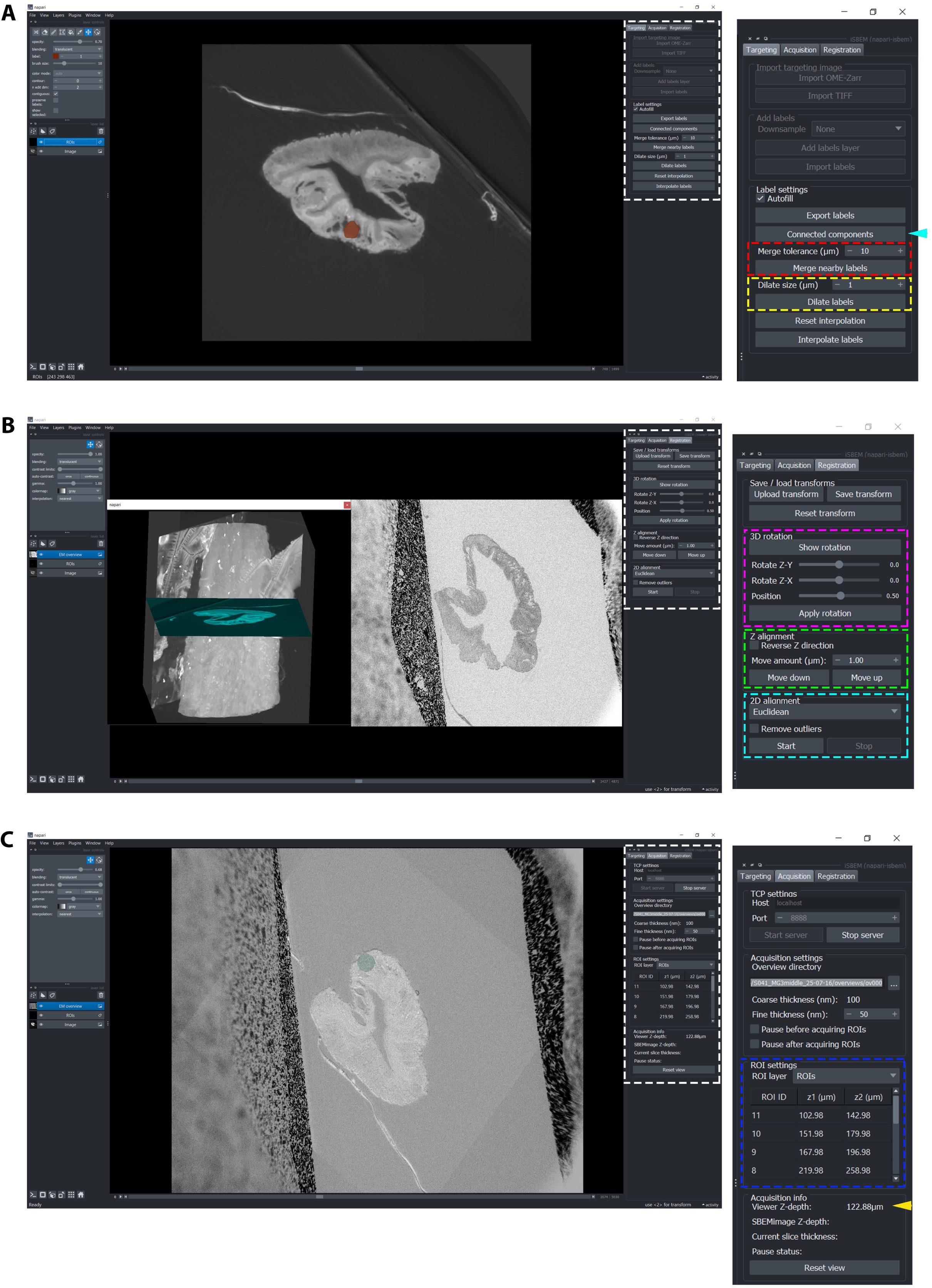
iSBEM graphical user interface. A. Targeting tab. XRM volume is imported (OME-ZARR (Moore et al., 2023) or TIFF format) and ROIs are either manually drawn using the napari segmentation functions (left panel) or imported as a binary tiff mask. An example is shown in brown in the visualization window. The right panel of the screen (white dashed box and further enlarged on the right) contains operations that can be done on the ROIs, such as dilation (yellow dashed box) and merging of nearby labels (dashed red box). The command “connected components” (cyan arrowhead) splits the imported binary masks into individual ROIs. B. Registration tab. Having acquired a few OV images and prompted iSBEM to their location, both XRM with ROIs and OV stacks are displayed in the visualization window. The registration tab allows registration of the XRM to the OVs. All transformation options are in the right panel (white dashed box and further enlarged on the right). The registration is done in 3 steps: 3D rotation, with the option to rotate the sample (XRM data) individually around single axes (dashed magenta box); Z alignment, with the option to reverse the Z direction, as in most cases the XRM stack is flipped with respect to the EM stack (green dashed box); 2D alignment allows a point-based registration in 2D, after the XRM volume has been rotated, with the possibility of choosing different transformations (Euclidean, similarity, affine), cyan dashed box. C. Acquisition tab. The right panel (white dashed box and further enlarged on the right) includes cutting thickness for low resolution imaging regimes (coarse thickness, which is read from the OV stack settings in SBEMimage) and high-resolution regime (fine thickness, which can be defined). After registration of the XRM volume in the “Registration” tab, the ROIs are assigned an ID number and their starting and ending depth in the SBF-SEM coordinate system are shown (blue dashed rectangle). At the bottom, further information regarding the run is shown, e.g. the depth of the image displayed in the visualization window (yellow arrowhead), or the current depth of the SBF-SEM stage.

### Registration of the XRM volume with the SBF-SEM coordinate system

When mounting a specimen onto the stage of the SBF-SEM, there is no flexibility to align it to a specific orientation. It is therefore of prime importance to precisely register the XRM 3D map of the sample with the actual resin block position in the microscope. For this, samples are inserted into the SBF-SEM and a few sections (one to ten) of the top-most part of the tissue are acquired at low resolution using SBEMimage overview (OV) settings. One of these OV images is then used to register the XRM data with the SBF-SEM coordinate system (Figure 2B). Using anatomical features visible in both modalities (eg. tissue boundaries, capillary lumen or nuclei, Figure S1), registration is achieved manually through three sequential steps: (i) a “3D rotation,” where a virtual slice of the XRM volume is rotated around the X and Y axes to match the OV orientation; (ii) a “Z alignment step,” where the rotated XRM slice is translated along the Z axis to match the OV height; and (iii) a “2D alignment,” consisting of point-based registration in 2D using different transformation types (Euclidean, similarity, or affine).

### Initiating automated acquisition

Once the XRM volume is registered to the SBF-SEM overview, ROI coordinates are transformed into the microscope stage coordinate system, allowing iSBEM to predict the Zstart and Zend positions of each ROI and to visualize them alongside the actual stage Z coordinate in the plugin (Figure 2C). These coordinates are then transferred to SBEMimage, where tiled acquisition grids spanning the full lateral extent of each ROI volume are pre-generated but held inactive. After each slice, iSBEM evaluates the current stage Z position against a virtual cross-section of the ROI segmentation mask at that depth and sends tile-level commands to SBEMimage for adaptive activation or deactivation of individual tiles within these grids (Figure 3J, Figure 4D-F, Figure S5B). This ensures that only tiles overlapping with the segmentation footprint at a given depth are acquired at high resolution, dynamically matching the acquisition footprint to the actual shape of the ROI through the Z axis. In the absence of any ROI segmentation at a given Z position, acquisition continues automatically with low-resolution overview images to maintain spatial context across the block. Independent cutting thicknesses can be specified for these two regimes, allowing thicker sections during low-resolution traversal and thinner sections during high-resolution ROI acquisition.

**Figure 3.**
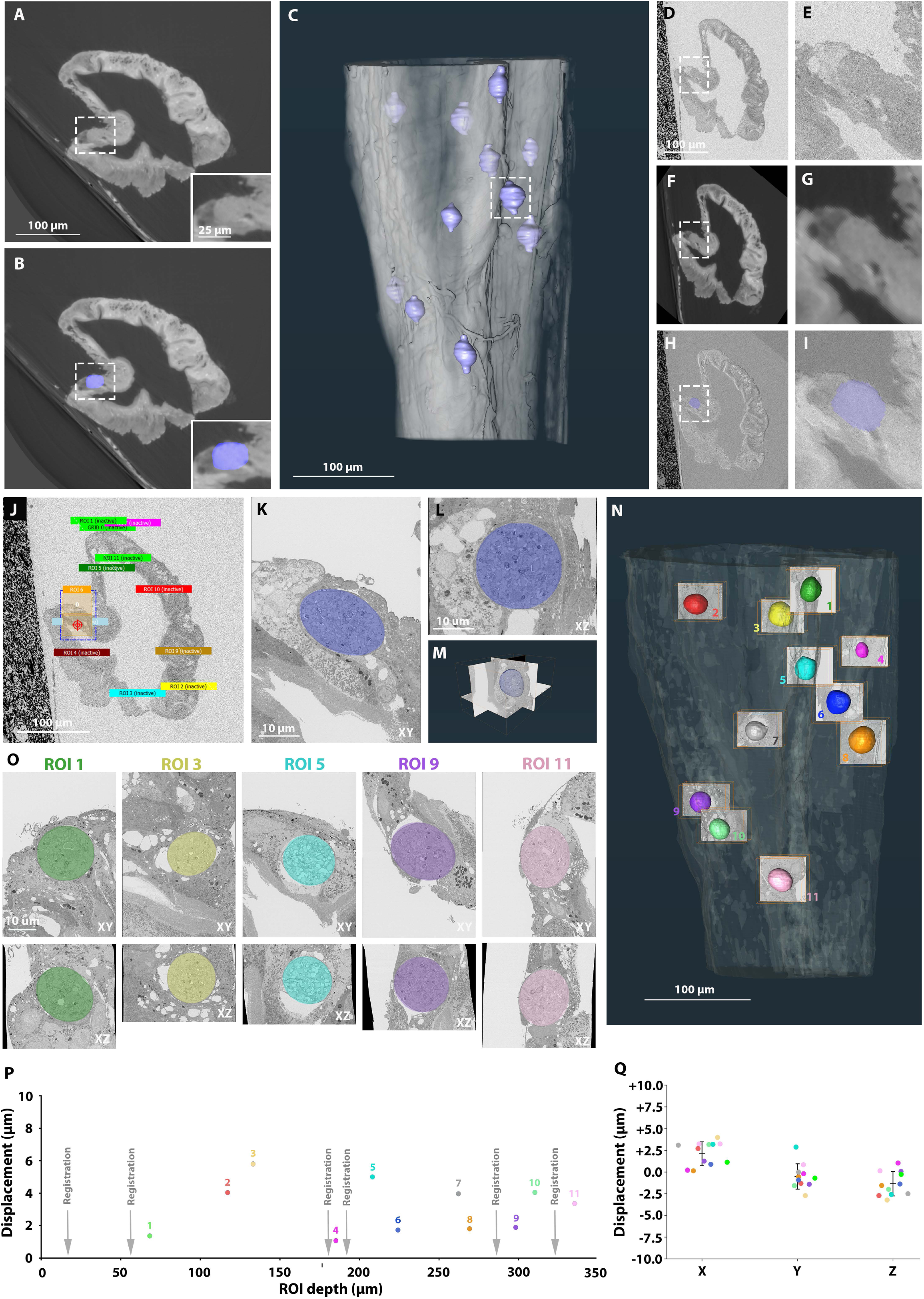
Semi-automated targeting of malarial oocysts in an infected mosquito midgut using iSBEM. A,B. Virtual slice of the XRM volume, showing a cross section of a malaria oocyst (dashed white box, enlarged in the inset). In B, the oocyst segmentation is overlaid in lavender. C. Volume rendering of the mosquito midgut (grey) from the XRM volume with the 11 segmented oocyst targets (lavender). To add a margin of error in Z for the acquisition, the segmentation was extended in Z, giving the round oocysts an elongated shape. The oocyst shown in A and B is highlighted by the dashed box. D,E. OV images displayed in iSBEM during the SBF-SEM acquisition. E is an enlargement of the boxed region in D. The highlighted oocysts are the same shown in A,B. F,G. Virtual slice of the XRM volume after registration, at the same location as in D,E. G is an enlargement of the boxed region in F. H,I. Overlay of the images in D,F and E,G, respectively, showing the precision of the registration. In lavender the oocyst segmentation. J. Snapshot from the SBEMimage interface, showing all the grids that were automatically generated from the segmented ROIs. ROI 6 is active at this point of the acquisition and the two tiles covering the oocyst are therefore highlighted in orange. K,L,M. High resolution imaging of ROI6, corresponding also to A,B,E,G,I. K shows a cross section of the oocyst (in blue) through its largest diameter in the XY plane. L shows an XZ view of the acquired volume and M the three orthogonal axes and a rendering of the oocyst in blue. The central location of the ROI in all the planes shows the precision of the targeting. N. Volume rendering of the mosquito midgut generated from SBF-SEM OVs (in grey transparent), with overlays of the acquired high resolution volumes for each of the 11 ROI. Shown is a XZ plane of the acquired and a rendering of the oocysts. All targets were entirely acquired, with minimal buffer volume. O. Central cross section of representative ROI volumes in XY (top) and XZ (bottom), with the targeted oocyst overlaid in colors. The central position of the oocysts in both planes shows the high precision of the targeting. P. Targeting accuracy measurement. The EM volume of each ROI was registered to the corresponding XRM crop around the segmented target (see Figure S2). The transformation vector is displayed in the plot (ROI displacement) as a function of the ROI depth. The grey arrows show the registration refinement points. Q. Displacement (euclidean distance) obtained for all ROIs in individual axes. Each dot represents an ROI, with colors matching those of the ROIs displayed in N.

**Figure 4.**
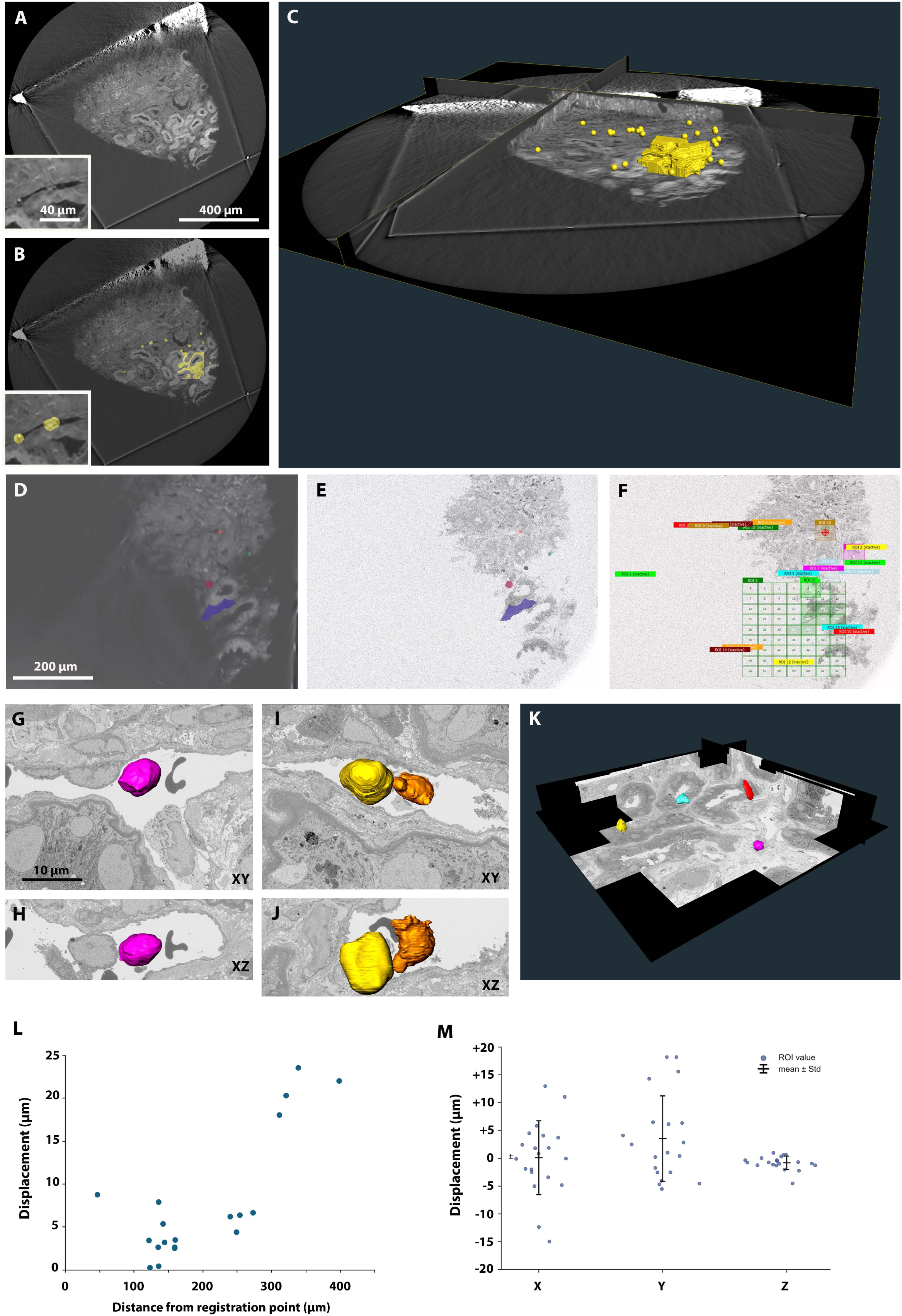
Fully automated targeting of immune cells in a human kidney biopsy. A. Virtual slice of the XRM volume, showing a cross section of a kidney biopsy. The inset shows putative immune cells identified in a peritubular capillary. B. Virtual slice of the XRM volume shown in A overlaid with a mask of the target ROIs (yellow). The small circles correspond to single immune cell targets. The larger convoluted region corresponds to peritubular capillaries (PTCs), our target region for identifying immune cells. C. 3D visualization of the biopsy slice with the small and large ROIs to be targeted by SBF-SEM in yellow. D. View of the iSBEM napari interface during SBF-SEM acquisition showing a virtual XRM slice with the ROIs that are exposed at this slice. E. View of the iSBEM napari interface during SBF-SEM acquisition showing the actual SBF OV image at the same depth as in D, with the prediction of the ROI location overlaid. Correspondence between the OV and the virtual XRM slice shows a good registration between the 2 volumes, which allows a precise targeting of the ROIs. F. View of the SBEMimage interface showing the grids and tiles that are activated by iSBEM on the section shown in D and E. G-H. Example of an intracapillary immune cell targeted with the iSBEM workflow, corresponding to the left yellow circle in B. Cell segmentation projected on an XY plane is shown in G and segmentation projected on the XZ plane in H. I-J. Example of 2 intracapillary immune cells targeted with the iSBEM workflow, corresponding to the fused right yellow circles in B. Cell segmentation projected on an XY plane is shown in I and segmentation projected on the XZ plane in J. K. 3D visualization of the EM volume of a large biopsy region covering peritubular capillaries (multi-tile grid seen in F), after tile stitching and slice alignment. Immune cells within peritubular capillaries are segmented and rendered in different colors. L. Displacement in 3D as a function of the distance from the initial registration point (see Materials and Methods). Each dot represents a single cell. Note the large offset for cells that have a distance > 300 µm from the registration point. M. Displacement (Euclidean distance) obtained for all single cell ROIs in individual axes. Each dot represents one ROI.

In principle, the acquisition can now run completely unsupervised until the maximum depth specified in SBEMimage is reached. However, depending on the initial registration accuracy, targeting may lose precision throughout the acquisition depth. We therefore recommend planning an acquisition pause after approximately 10 µm depth to assess registration accuracy (Figure 1C). This can be evaluated in iSBEM by inspecting the overlay between the last OV and the predicted corresponding virtual XRM slice. Registration can then be refined, or acquisition can be resumed. In our experience, registration accuracy varies depending on the abundance of anatomical features visible in both XRM and low resolution OVs in the first portions of the sample (Figure S1). For critical samples, we advise visual monitoring once daily with refinement when necessary.

### Acquisition of malarial oocysts in mosquito midguts

As proof of concept for the workflow, we targeted *Plasmodium* oocysts in infected mosquito midguts. These stages of the malaria parasite produce *Plasmodium* sporozoites, the mosquito transmitted forms. After dissection of midguts and EM sample preparation, resin embedded samples were mounted on a pin stub with the midgut’s longest axis aligned vertically, perpendicular to the stub surface, resulting in tissue approximately 400 µm long in the Z direction. The samples were scanned by HiTT, where oocysts appeared as spherical objects with characteristic texture embedded within the surrounding tissue (Figure 3A).

We selected 11 to 15 oocysts with broad depth distribution throughout the volume and segmented them manually using Amira (Thermo Fisher Scientific) or the napari segmentation tools available through iSBEM (Figure 3B,C, S3A, S4A). To provide a buffer in Z, segmentation was extended by 2–5 µm in the Z direction above and below the estimated oocyst edges (Figure 3C, S3A, S4A). Samples were then mounted onto the SBF-SEM stage and processed through the iSBEM workflow. Figure 3D–I shows an example overlay between the EM OV, the registered XRM virtual slice, and the transformed segmentation of an oocyst, demonstrating accurate registration. Acquisition of the oocysts was then performed automatically with SBEMimage, triggered by iSBEM. Figure 3J shows the activated tiles covering the ROI displayed in Figure 3A–I. To assess targeting precision, we display an XY view (Figure 3K) and XZ view (Figure 3L) of the oocyst at its largest cross-section, as well as a 3D rendering (Figure 3M) of the complete acquired ROI volume. Measuring the distance between the oocyst center and the image center provides the targeting accuracy. In this example, the measured distance was 1.7 µm.

In the experiment shown in Figure 3, we targeted and successfully imaged 11 ROIs at a 10×10×50 nm^3^ voxel size (Figure 3N). The run was continuously monitored, checking the quality of the registration between the latest acquired OV and the predicted XRM XY plane. When we observed a significant offset between the two imaging modalities, we paused the run to refine the registration manually. This happened 6 times (a 5 to 10 minutes intervention) over the 8-day long acquisition in this example (Figure 3P). All oocysts were fully acquired with minimal non-useful imaging time (i.e., time spent acquiring non-oocyst tissue at high resolution). To estimate the gain in acquisition throughput, we calculated the time that would be required to image columns spanning the full depth of the tissue that would capture all the chosen ROIs. We consider this sham imaging regime would be the closest scenario to a CXEM experiment targeting multiple ROIs while aiming to minimize human intervention but lacking the support of iSBEM (see methods). In this example, the targeted acquisition scheme would have taken ∼ 75 days. Compared to the 7.5 effectively achieved with iSBEM, the throughput gain is estimated to be 9.9 fold. While the acceleration factor is sample dependent and is determined by total tissue size, target frequency and distribution, the concept of throughput gain can be generalized.

We also assessed targeting precision by registering the XRM volume cropped around each ROI with the corresponding EM volume and extracting the displacement vector of the transformation (Figure S2; see also Materials and Methods). Thanks to the XRM-to-OV registration correction along the run, the accuracy in the 3D space was kept below 5.8 µm (Figure 3P). Displacement around the individual axes was always below 4 µm (Figure 3Q).

We then investigated whether acquisition could run unsupervised. To this end, two additional infected mosquito midgut samples were prepared and imaged identically to the one described above, with the exception that, after initial registration and further refinement at 10 µm acquisition depth, the runs were left unsupervised for the remaining ∼350 µm. As expected, experiment success depended on initial registration precision. With precise initial registration, acquisition was nearly as successful as with the supervised strategy (Figure S3) with a maximum 3D displacement of 6.01 µm. In this case, we acquired 100% of 10 out of 11 oocysts, with only a small fraction (∼1-2%) missing of the 11th oocyst (Figure S3C, ROI4).

However, when insufficient registration landmarks within the first portion of the sample prevented precise initial registration, targeting accuracy degraded significantly with acquisition depth (Figure S4). Even in such a case, we acquired the first 9 oocysts entirely (Figure S4C). However, from a depth of ∼220 µm onwards, the error became too large (up to 20.68 µm, Figure S4D) and considerable fractions of the last 2 targets were missed (estimated 20 to 40%). Note that the largest targeting offset was measured in the x and y axes (up to 17.7 µm), and less prominent in z (up to 4.8 µm) as expected for samples of a vertical geometry (Figure S4E).

### Acquisition of intracapillary immune cells in human kidney biopsies

The infected mosquito midguts were mounted vertically on the stub, resulting in samples elongated primarily in the Z direction (depth) with relatively small XY target spread (maximum midgut cross-section width was ∼300 µm). To test the workflow with samples having more XY-distributed targets, we used vibratome slices of two human kidney biopsies (Burrell et al., 2025) and aimed to acquire intracapillary immune cells. These samples were slices of ∼50 µm thickness with an approximate area of 900 × 700 µm^2^, with targets distributed throughout the entire tissue.

Our strategy was to target both individual putative immune cells identified by XRM as well as a larger biopsy volume containing both glomerular capillaries and peritubular capillaries (PTC) where immune cells are potentially enriched (Figure 4A–C, Figure S5A). We identified 24 putative intracapillary single immune cells in the first sample analyzed (Figure 4) and 15 in the second (Figure S5), distributed in glomerular capillaries or PTCs. These were manually identified and spheres with a diameter of 14 µm (Figure 4) or 22 µm (Figure S5) were used as masks for further targeting. When cells were in close proximity (< 10 µm), ROIs were merged using iSBEM’s “merge nearby labels” function (Figure 2A). As a result, the experiment was run for 20 and 13 targets, respectively. To further test the versatility of iSBEM for defining ROIs, we created a larger, convoluted target that captured the lumens of multiple adjacent PTC where immune cells are usually found. Note that in the example shown in figure 4F, this convoluted ROI was imaged with different tile size compared to the other 13 ROIs. This illustrates the ability of the iSBEM / SBEMimage workflow to adapt imaging parameters to each individual target.

For the first two experiments shown here (Figure 4 and Figure S5), single immune cells were covered by individual tiles measuring 40 × 30 µm^2^(Figure 4G-J, S5C). The number of slices was determined by the sphere diameter (14 or 22 µm) and the sectioning thickness (50 nm). The segmented PTC lumens triggered the creation of a larger grid made of 63 tiles each measuring 30×23 µm^2^ (Figure 4F,K), or 30 tiles each measuring 40×30 µm^2^ (Figures S5B,D). As described above, high-resolution acquisition is driven by adaptive activation of individual tiles based on the overlap between the segmented ROI (Figure 4D and E) and the SBEMimage grid at each Z plane, ensuring that only tiles corresponding to the segmented object footprint are acquired (Figure 4F).

Unlike the mosquito midguts, which had a significant depth (>300 µm), the processed kidney slices were only ∼50 µm thick. We thus opted to run these acquisitions in an unsupervised manner. However, because the slices were not mounted completely parallel to the stub, the tissue was exposed obliquely, resulting in only a corner of the tissue being imaged at the beginning of the SBF-SEM experiment. This often provided insufficient anatomical features for precise XRM-OV registration (Figure S1). To overcome this problem, it was particularly important to plan a second registration after a few microns of acquisition. We noticed that within 10 µm depth, targets could still be safely acquired even without correction. At this point though, registration had to be inspected and modified before resuming the unsupervised acquisition of the remaining tissue depth. Although during these experiments we acquired both single cells and larger multi-tile volumes, the evaluation of the targeting accuracy was limited to the single cell ROIs. In the analysis of these samples, which have a horizontal geometry, we did not observe a dependency of targeting precision on target depth; rather, the distance of the ROI in XY from the area where the registration was initially performed seemed to play a role. In one of the samples (Figure 4) the registration between EM OVs and XRM volumes was performed on the first exposed corner of the sample, and the largest offset in the XY plane (up to 23.6 µm) were observed in ROIs located in the opposite corner of the sample, at a 300 µm lateral distance from the registration point (Figure 4L,M). For the second sample though, the targeting offset stayed below 12.2 µm (Figure S5E,F), even if the initial registration was performed only in one corner of the biopsy. This variability in targeting accuracy shows once more the importance of the registration step. The more visible cues are available for the registration, the more precise the targeting will be.

Even with large targeting offsets in XY, the dimensions of the tiles (40×30 µm^2^) enabled capturing most of the targets at their full volume (Figure 4G-J, Figure S5C). Considering both experiments, 27 out of 33 targets were fully acquired (82%), 5 were partially covered (15%), and 1 was completely missed (3%).

Although targeting was largely successful, analysis of acquired EM volumes revealed that not all targeted cells were immune cells (Figure S6) as initially interpreted when inspecting the XRM data. In several cases (12, corresponding to 36%), immune cell prediction was incorrect, as endothelial cell soma bulging into the capillary lumen can appear very similar to circulating blood cells at HiTT resolution. To improve correct immune cell identification, we implemented a correlative light, X-ray, and electron microscopy approach (CLXEM). We employed the complete nanopathology pipeline described in (Burrell et al., 2025). Before EM sample preparation, biopsy slices were immunolabeled with an antibody against the extracellular CD16 epitope, which worked on tissue without permeabilization, and imaged by confocal microscopy. After sample preparation for EM, kidney slices were imaged by HiTT. LM data were then overlaid with XRM volumes, and the combination of both imaging modalities was used to define ROIs for the iSBEM pipeline. For this experiment, we selected 25 single cells within the sample that were either CD16-positive or CD16-negative yet unambiguous immune cells visible in the capillary lumen. In this example, we designed precise segmentation of target cells instead of spheres (Figure 5A-D).

**Figure 5.**
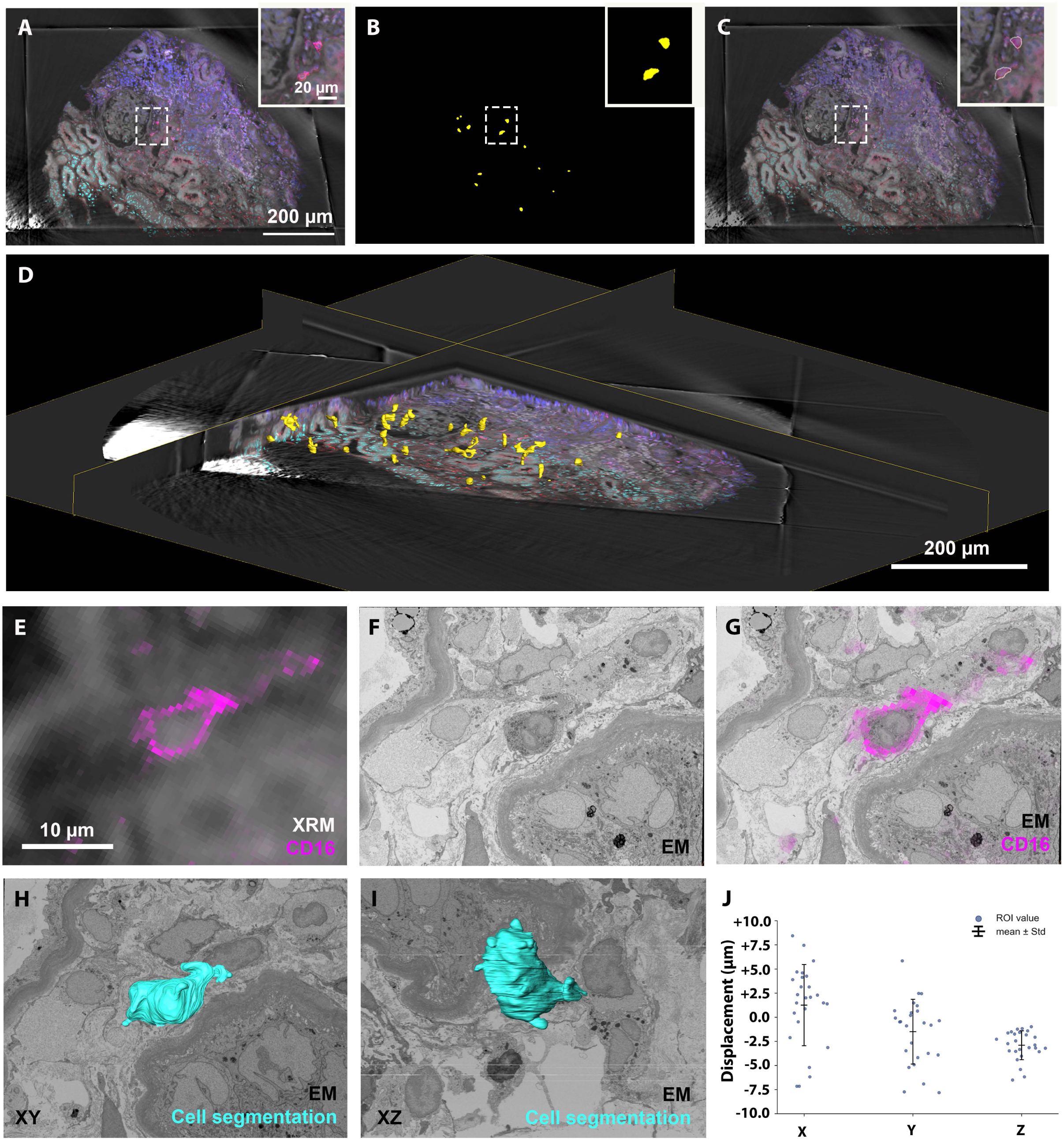
Automated targeting of immune cells in a human kidney biopsy based on correlative light and X-ray imaging. A. Pre-embedding immunofluorescence stack overlaid with post-embedding XRM volume. The biopsy slice was imaged from both sides and different colors were assigned to each side. Blue and cyan: Hoechst; Red and magenta: CD16. XRM is shown in greyscale. The inset shows an enlargement of the dashed box area. B. Masks of the immune cells of interest, identified by morphological criteria and CD16 expression. C. Overlay of the image in A with contours of the immune cell masks (in yellow). D. 3D visualization of the correlative LM-XRM dataset, with segmentation of the target cells in yellow. E-G. CLXEM overlays. The 3 imaging modalities have been registered. E shows the overlay of pre-embedding LM and post-embedding XRM in the region of a CD16 positive cell, which was one of the acquisition targets. F shows the high-resolution EM image of the targeted immune cell. G shows overlay of the LM data with EM. H-I. Segmentation of the cell shown in E-G. XY view in H and XZ in I. J. Displacement (Euclidean distance) obtained for all ROIs in individual axes. Each dot represents an ROI.

To provide a margin of error for automated acquisition, cell masks were dilated in 3D by 2 µm in iSBEM using the “dilate size” command in the napari plugin (Figure 2A). Due to their convoluted shapes, some cells fit within a single tile (40 × 30 µm^2^) while others required coverage by 2 or 4 tiles. With an unsupervised approach, we completely acquired 20 out of 25 targets, consistent with previous acquisition results. The remaining 5 (20%) were partially covered thus no targets were completely missed in this experiment, achieving a success rate similar to other kidney biopsy experiments. However, due to the correlative LM and XRM volumes used for target identification, 100% of targets in this experiment were immune cells. This improved the fraction of fully acquired correct targets from 54.5% when only XRM is used for targeting to 80% when CLXEM is employed.

To evaluate the gain in throughput for such experiments, we compared the time spent to acquire the 25 cells using iSBEM with the estimated time that would be required to acquire at high resolution the ROI-containing tissue columns. With iSBEM, the imaging time for 25 cells at 10 x 10 x 50 nm^3^ was 174 hours. At the same pixel dwell time and scale parameters, imaging the corresponding columns would have taken 517 hours. The gain in this experiment was then 3 fold.

## Discussion

SBF-SEM is suited for sub-cellular imaging of large volumes. It has been used to acquire large scale tissues or even entire organisms. However, frequently, to answer a biological question, it is sufficient to image small volumes within a large specimen. In such cases, targeting the acquisition to defined ROIs is critical, as it allows reduction of imaging time and increased throughput. To this aim, correlative X-ray and electron microscopy (CXEM) has shown great potential because it allows visualization of anatomical features that can be used as registration landmarks. However, most workflows that employ XRM to target vEM are manual and based on comparing XRM virtual slices and toluidine blue stained sections imaged by light microscopy, or low resolution OVs from the SBF-SEM. These require either continuous interaction or very close supervision during the acquisition, making it very challenging and time consuming to target many ROIs. Our work aims to fill this gap and provides open access software tools as well as an easy workflow to target multiple ROIs in large resin embedded samples in a fully or partially unsupervised way.

The workflow makes use of iSBEM, an original napari open-source plugin, which provides tools for data import (ome.zarr and tiff data formats available), basic segmentation, multimodal volume registration in 3D, connection to SBEMimage to drive image acquisition at the ROIs and constantly updated visualization of the overlay between the predicted current virtual slice and the actual OV for registration inspection. Using these tools, we were able to automatically acquire volumes alternating a low-resolution regime for slices with no ROI present (i.e. OV acquisition and cutting thickness of 100nm) and high resolution only where ROIs were present (i.e. OV acquisition plus high resolution of 30 x 40 μm^2^ tiles covering the ROI location and a cutting thickness of 50 nm). We opted to acquire low resolution OVs throughout the sample, to provide a spatial context for the ROIs. This was also important to perform debris detection after each cut, with a minor imaging time overhead (∼ 10 s per slice).

A core concept of iSBEM is the acquisition of selected sub volumes of interest at high resolution (CXEM targeting). To evaluate CXEM throughput gain, we thus estimated the time that would have been necessary for imaging the same targets whilst trying to minimize human intervention, and compared it to the observed experimental time. Across the samples analyzed we observed a throughput increase ranging from 3 to 10-fold.

To better appreciate iSBEM efficiency, it is appropriate to compare the workflow with a standard CXEM experiment. Manual CXEM targeting requires the operator to identify each ROI in real time on successive SBF-SEM overviews, configure the high-resolution acquisition window, and monitor imaging before proceeding to the next ROI. Because SBF-SEM runs typically span several days, this serial mode of operation ties the operator to the microscope for the duration of the experiment and effectively caps the number of ROIs that can realistically be acquired in a single run. The burden compounds when targets are numerous or distributed irregularly through a large depth: each ROI must be re-located independently from a limited set of anatomical landmarks, and reproducing precise targeting decisions consistently across days of acquisition is both impractical and operator-dependent. iSBEM removes these constraints by registering all ROIs once on the XRM map and queueing them for unattended acquisition, decoupling experiment duration from operator availability and yielding deterministic, reproducible targeting regardless of ROI number or geometry.

During the development of the workflow, we used two kinds of samples to test the performance with differently distributed targets. For the *Plasmodium* oocysts, because of the elongated shape of the mosquito midguts and their vertical orientation in the resin block, the ROIs had a relatively small lateral distribution, but were spread over a large depth (up to almost 400 μm). On the other hand, the kidney vibratome slices were much thinner (50-60 μm), but the planar distribution in X-Y was over almost a millimeter. As expected, in all cases the targeting precision was very good in proximity to the region where we performed the registration between XRM and the first OVs of the EM acquisition. When we ran the acquisition completely unsupervised, we noticed that the precision tended to decrease, mostly because of small inaccuracies in the rotation of the XRM volume, which had a big effect at distance from the registration point. Such inaccuracies originate most probably in the resolution gap between the 2 imaging modalities. XRM data are acquired at 650 n m isovoxel size, while SBF-SEM2D OV images, used for the registration, are typically between 100×100 to 250×250 nm^2^ in XY and display information from the surface of the exposed block. As a consequence, the registration can appear adequate for multiple slight tilt angles of the XRM XY plane at a given Z level. This resulted in a bigger inaccuracy for the deepest oocysts and for the immune cells localized in the corner of the biopsy opposite to the registration site, in some unsupervised acquisitions (e.g. Figure S4, Figure 4). In some instances, the initial registration proved good enough to allow for a precise targeting throughout the sample even in fully unsupervised acquisitions (e.g. Fig. S3, S5). In such unsupervised cases, it would nonetheless be good practice to design large acquisition tiles to compensate for slight registration inaccuracies.

Further development of the workflow could use a recurrent automated registration check and eventually re-registration if needed. However, with the current version of the software, we recommend to visually inspect the registration accuracy during acquisition, e.g. once or twice per day. iSBEM has an interface to monitor the registration where each newly acquired OV image is automatically loaded to the stack that is overlayed with the predicted current virtual slice of the XRM volume, making the check quick and easy. If the registration is not precise enough, the acquisition could be paused and the registration refined, a process that normally takes ∼5 to 10 minutes. Such a supervised mode of acquisition thus does not impact significantly the overall throughput of the experiment.

In conclusion, the iSBEM workflow is an open-source solution that can be implemented in many EM labs, facilitating and streamlining the targeted acquisition of diverse biological samples.

## Material and methods

### Mosquito dissection and preparation for SBF-SEM

At 9 days post infection with *Plasmodium berghei*, *Anopheles stephensi* mosquitoes were transferred into 15 mL tubes, which were placed on ice to induce immobilization. Immobilized mosquitoes were briefly submerged in 100% ethanol and subsequently transferred into PBS, where they remained until dissection. Midguts were dissected under a dissecting microscope using 1 mL syringes fitted with sharpened needles. Dissection was initiated by piercing the thorax with one needle while gently pulling the terminal abdominal segments with the other to extract the abdominal organs. The midgut was isolated by making incisions at the lower esophagus and at the posterior end of the gut. Samples were immediately transferred to the primary fixative (2.5% (v/v) glutaraldehyde (EMS), 2% (v/v) paraformaldehyde (EMS) and 5% sucrose (wt/v)) in 0.1 M cacodylate buffer) until further processing (∼2h). The following steps were performed using a Pelco BioWave Pro + (Ted Pella, Inc.), similarly to Darif et al., 2026. Each microwave incubation was done with 2 min on/off microwave cycles at 100 W under vacuum, for a total of 14 min, unless otherwise specified. After the first fixation following dissection, samples underwent another fixation in the Biowave in a solution containing 1.5% glutaraldehyde, 2.5% paraformaldehyde, and 5% sucrose in 0.1 M cacodylate buffer. Following fixation, samples were rinsed with 0.1 M cacodylate buffer (one immediate wash, followed by two washes of 40 s each). Post-fixation was performed in 2% osmium tetroxide in 0.1 M cacodylate buffer in the Biowave, followed by osmium reduction with 2.5% potassium ferrocyanide (K₄[Fe(CN)₆]·3H₂O) in 0.1 M cacodylate buffer in the Biowave. Subsequent washes were performed with distilled water (one immediate wash, followed by two washes of 40 s each in the microwave). Samples were then incubated in 1% thiocarbohydrazide (Sigma Aldrich) aqueous solution in the Biowave, followed by additional water washes as described above. A second osmium staining step was performed using 2% aqueous osmium tetroxide in the Biowave. Dehydration was carried out through a graded ethanol series (25%, 50%, 75%, and three changes of 100%), each for 40 s at 250 W without vacuum. Samples were then infiltrated with increasing concentrations of Epon (Serva) resin (25%, 50%, 75%, 90%, and three changes of 100%), each for 3 min at 150 W under vacuum. Final embedding in 100% Epon was performed overnight at room temperature. Polymerisation was completed by incubating samples at 60 °C for 48 h.

### Preparation of human kidney biopsy for SBF-SEM

Sample collection and preparation was performed as detailed in Burrell et al., 2025. Material excess to diagnostic needs from indication kidney transplant biopsy samples were taken at Imperial College NHS Trust Renal Unit. Imperial College Healthcare NHS Trust Tissue Bank has ethical approval to both collect human tissue excess to diagnostic needs and release material to researchers (MREC 22/WA/2836). Patients undergoing a procedure at Imperial College Healthcare NHS Trust were asked at time of the procedure if they consented to material surplus to diagnostic needs from their tissues being used for research, using patient information sheet (PIS) v.8. Patient responses were recorded in the electronic patient record and on the pathology request form. Approval for our specific project was registered with Imperial College Healthcare Tissue Bank as project R14094: Outcome analysis after renal transplantation in patients with de novo donor-specific antibodies.

The biopsy tissues were fixed in 4% EM-grade formaldehyde (FA) (TAAB) in 0.1M phosphate buffer (PB) overnight at room temperature and then stored in 1% FA in 0.1M PB at 4°C until further processing. Next, samples were vibratome sectioned after embedding in 3% low melting point agarose (Sigma Aldrich) in 0.1M PB. The 60 μm thick slices were further stored in 1% FA in 0.1M PB at 4°C. The sample shown in Figure 5 underwent an additional immunofluorescence labeling as described in Burrell et al., 2025. Preparation for SBF-SEM was performed using a modified version of the protocol from (Deerinck et al., 2010), as detailed in Burrell et al., 2025.

### X-ray imaging (XRM)

High-throughput X-ray imaging (HiTT) was conducted at the EMBL beamline P14 of the PETRA III storage ring (DESY, Hamburg, Germany). Sample blocks were mounted on an MD3 diffractometer using magnetic SPINE-style goniometer bases. Resin-embedded samples were either trimmed and fixed to metal pins using acrylate glue or mounted onto custom-made 3D-printed bases attached to the goniometer bases. Automated sample mounting was performed with the MultiAxesRoboticVersatileINstaller (MARVIN) system.

Imaging was performed at X-ray energies ranging from 12 to 23 keV. Further details of the beamline setup, image acquisition, and processing are described in Albers et al. (2024) and Burrell et al., 2025. Image acquisition, flat-field correction, and reconstruction were automated. Voxel sizes were 650 nm with the 10× objective and 325 nm with the 20× objective.

### Segmentation of ROIs for the iSBEM pipeline

#### *Plasmodium* oocysts in the mosquito midguts

The HiTT volume was imported in Amira (version 2023, Thermo Fisher Scientific) and a fraction of the oocysts present in the midgut were manually segmented with the paint brush tool and interpolation. Alternatively, the ROIs were directly segmented after XRM data import using the iSBEM napari interface. The targets were intentionally chosen with a large spread over the volume to test the iSBEM pipeline with ROIs that reach a large depth in the resin embedded samples. To introduce a margin of error for the imaging workflow, the segmentation was extended in Z beyond the ends of the oocysts, resulting in a spheroid shape.

#### Immune cells in the kidney biopsies

For the CXEM experiments (Figure 4 and S5), the HiTT volumes were inspected and potential intracapillary immune cells were identified by morphology, density, and position in the capillaries. The potential single cell targets were segmented using the sphere tool in IMOD (Kremer et al., 1996), with a fixed diameter of 14 and 22 μm for the experiments shown in Figure 4 and S5, respectively. For the larger PTC regions, the ROIs were manually drawn to cover regions with high capillary density around the renal tubules.

For the CLXEM experiment (Figure 5) the targets were identified using an overlay of the HiTT and immunofluorescence data. The fluorescence images were produced as described in Burrell et al., 2025. Briefly, before EM sample preparation the vibratome slices were immunolabelled for CD16 without permeabilization. Hoechst was used as a counterstain for the nuclei. Imaging was then performed with a confocal microscope (Zeiss LSM900, equipped with a 40X 1.2 NA water immersion lens) from both sides for full high quality volume imaging of the entire slices. As targets for iSBEM we selected both CD16 positive cells and CD16 negative intracapillary cells (that had a high probability of being immune cells). The targets were manually segmented in IMOD, precisely drawing the contours of the cells.

### iSBEM

iSBEM is a plugin for the Python-based interactive image viewer napari. It provides real-time visualisation and interactive control of SBF-SEM acquisitions via the separate, open-source acquisition software SBEMimage. The plugin uses napari’s interactive multi-dimensional image viewer to enable simultaneous visualisation of XRM and SEM data. 3D ROIs in the XRM coordinate system can be defined within the plugin or imported as a labelled TIFF mask and are represented in napari as a labels layer. User-guided registration between the two modalities produces a transformation matrix that, when combined with pixel size metadata, enables conversion of these ROIs to physical SBF-SEM stage coordinates.

iSBEM and SBEMimage communicate through two distinct methods: file monitoring and transmission control protocol (TCP) socket communication. The plugin continuously monitors the SBEMimage output directory and dynamically loads overview slices at intervals matching the defined coarse cutting thickness, building a 3D volume of the ongoing acquisition. Through the TCP connection, SBEMimage sends a status update to the plugin after each slice acquisition, including acquisition state and current stage z-height. The plugin responds with commands to add grids, activate or deactivate grid tiles, adjust cutting thickness, or pause acquisition, enabling automated acquisition of the user-defined ROIs.

The iSBEM napari plugin is released under the MIT software license and is available via GitHUb at https://github.com/FrancisCrickInstitute/napari-isbem. Detailed user instructions and documentation are available in the GitHub repository and in the documentation at https://franciscrickinstitute.github.io/napari-isbem. The plugin was developed using napari version 0.5 and the dev branch of SBEMimage. The results presented in this study were obtained using napari-isbem version v0.1.0.

### Analysis of the targeting precision

Targeting accuracy was determined by performing rigid registration between the ROIs in the X-ray scan and the corresponding high-resolution EM images. The spatial offset between the initial X-ray scan and the acquired EM images was used as a quantitative estimate of targeting precision (see Figure S2).

#### HiTT data pre-processing

To be able to apply the rigid registration, the respective ROIs were cropped from the X-ray scan based on the center of the binary mask used to define ROIs, to match the size of the high-resolution EM image dimensions (40 µm x 30 µm). Prior to cropping, the rotation and affine transformations used during registration were applied to the X-ray scan. After cropping, each image was manually assigned the corresponding ROI identity and renamed accordingly. Following ROI assignment, X-ray images were pre-processed by inverting the image LUT. Subsequently, the appropriate output directory was generated and the pre-processed X-ray images were moved to the respective output directory.

#### EM data pre-processing

EM images were downscaled to match the voxel size and slice number of the X-ray images using averaging and bilinear interpolation, and then a Gaussian blur algorithm was applied (σ = 1.6). Ultimately, the minimum display value was set to 100 to match the contrast of the X-ray tomograms.

#### Registration and displacement measurement

Rigid registration between pre-processed HiTT and EM volumes was performed using the ITK-Elastix package (Klein et al., 2010) with the default ParameterMap, where “AutomaticTransformInitialization” was set to false, the fixed and moving image dimensions were set to 3, and the maximum number of iterations was set to 1000 and the maximum step length was set to 1.0. Furthermore, the Adaptive stochastic gradient descent optimizer and Advanced Mattes Mutual Information were used. For image sampling, “RandomCoordinate” was used, with the maximum number of sampling attempts set to 32, the number of spatial samples to 2000 and retrieving a new sample every iteration was set to true. A multi-resolution pyramid strategy was implemented with 5 resolutions and a pyramid schedule ranging from σ = 8 to 4, 2, 1 and finally 0. Lastly, a linear interpolator was used. The registration results were visually inspected, and the offsets of successfully registered ROIs were used to plot the average displacement as a measure of targeting precision.

#### Plots representing displacement as a function of depth

(Figures 3P, S3E and S4D). The displacement measurements calculated as described above are plotted against the acquisition depth of each oocyst, defined as their starting Z coordinate.

#### Plots representing displacement as a function of distance from the registration point

(Figures 4L and S5E). The registration point was defined as the center of the area that was exposed on the block surface and was used for the registration at the beginning of the acquisition. The distance of each ROI was measured as a segment connecting the registration point to the center of the ROI sphere in the XRM dataset.

### Estimation of throughput gain with iSBEM

The imaging time to acquire CXEM data using iSBEM was experimentally determined considering the first and last OV image acquired, to which we subtracted the idle time, due to machine errors or pauses in the acquisition. This time therefore includes OV acquisitions, tile acquisitions, cutting time and sweep movements and OV reimaging, in case of debris detection.

The time necessary to acquire ROIs-containing columns was estimated using SBEMimage, after importing the ROIs from iSBEM, activating the tiles and defining the depth required to cover all the ROIs in the volume. This estimation does not take into account the debris detection workflow and is therefore an underestimation of the real time that would take to acquire the corresponding volume experimentally. Our estimations are therefor likely an underestimation of the actual throughput gain.

### Tile stitching and slice alignments

Single-tile ROIs were aligned in Z using the Fiji plugin “linear stack alignment with SIFT” using translation as transformation.

Up to 4-tile ROIs were stitched and aligned with TrackEM2, using translation for both transformations.

The larger volumes were aligned using Webknossos with Voxelytics (scalable minds, Potsdam, Germany). Voxelytics was configured to perform a least-squares optimization of SIFT (Lowe, 2004) feature matches, minimizing match distances between neighboring tiles in the 3D tile grid. False matches were excluded using a RANSAC approach (Fischler and Bolles, 1981). The optimization was performed in two steps: first solving only for translations, followed by solving for one affine transformation per tile with regularization.

### Segmentation and volume visualization

Features of the EM high resolution volumes of malarial oocysts were manually segmented using the paint brush tool and interpolation in Amira (Thermo Fisher Scientific). Registration of the high resolution SBF volumes to the low resolution OVs was performed in Amira using the manual volume editor.

Segmentation of the immune cells from kidney biopsies was done in Webknossos (scalable minds, Potsdam, Germany) using the quick select tool, followed by manual correction where needed.

Visualization of the segmented data was done with Amira.

## Code availability

### iSBEM

This napari plugin can be accessed from the Napari Plugin Hub, searching for napari-isbem, selecting version v0.2.0. Please refer to the official napari documentation on installation and use including installation of plugins. Detailed user instructions and documentation are available in the GitHub repository and in the documentation at https://franciscrickinstitute.github.io/napari-isbem.

### SBEMimage

This can be accessed from github, by selecting release 2026.06 from: https://github.com/SBEMimage/SBEMimage/. User documentation including installation instructions can be found here: https://sbemimage.github.io/SBEMimage.

## Acknowledgements

Human samples used in this research project were obtained from the Imperial College Healthcare Tissue Bank (ICHTB). ICHTB is supported by the National Institute for Health Research (NIHR) Biomedical Research Centre based at Imperial College Healthcare NHS Trust and Imperial College London. ICHTB is approved by Wales REC3 to release human material for research (22/WA/2836). EMBL ethical approval for the Schwab/ Duke/ Uhlmann project: A Nanopathology Platform for the prediction and early detection of disease in kidney transplant rejection (BIAC 2022-032) was granted in December 2022.

We acknowledge funding for staff and resources from the UKRI Medical Research Council (grant number MR/W031426/1). This work was supported by the Francis Crick Institute which receives its core funding from Cancer Research UK (CC1076), the UK Medical Research Council (CC1076), and the Wellcome Trust (CC1076). Candice Roufosse is supported by the NIHR Imperial Biomedical Research Centre (BRC). We acknowledge funding for staff and resources from the European Molecular Biology Laboratory. Part of this work was funded by the Deutsche Forschungsgemeinschaft (DFG, German Research Foundation) –Project number 240245660 – SFB 1129 (project Z2).

We would like to particularly thank the Electron Microscopy Core Facility staff at the EMBL for the strong support and advice; Christian Tischler and Julian Hennies (EMBL) for their input on the analysis of target precision; Mandy Boermel, Nadav Scher and Davi Bock for critical feedback on the manuscript; Adam McLean, Imperial College Healthcare Trust, Renal Unit, and Linda Moran, Electron Microscopy Unit, North West London Pathology, Charing Cross Hospital for support in handling of human biopsy samples.

## References

Albers, J., M. Nikolova, A. Svetlove, N. Darif, M.J. Lawson, T.R. Schneider, Y. Schwab, G. Bourenkov, and E. Duke. 2024. High Throughput Tomography (HiTT) on EMBL beamline P14 on PETRA III. J. Synchrotron Radiat. 31:186–194. doi:10.1107/s160057752300944x.

Bishop, D., I. Nikić, M. Brinkoetter, S. Knecht, S. Potz, M. Kerschensteiner, and T. Misgeld. 2011. Near-infrared branding efficiently correlates light and electron microscopy. Nat. Methods. 8:568–572. doi:10.1038/NMETH.1622.

Bosch, C., T. Ackels, A. Pacureanu, Y. Zhang, C.J. Peddie, M. Berning, N. Rzepka, M.-C. Zdora, I. Whiteley, M. Storm, A. Bonnin, C. Rau, T. Margrie, L. Collinson, and A.T. Schaefer. 2022. Functional and multiscale 3D structural investigation of brain tissue through correlative in vivo physiology, synchrotron microtomography and volume electron microscopy. Nat. Commun. 13:2923. doi:10.1038/s41467-022-30199-6.

Briggman, K.L., and D.D. Bock. 2012. Volume electron microscopy for neuronal circuit reconstruction. Curr. Opin. Neurobiol. 22:154–161. doi:10.1016/J.CONB.2011.10.022.

Burrell, A., M. Lawson, P. Ronchi, X. Xiong, J. Albers, A. McLean, M. Willicombe, L. Moran, J. Hounsome, M. Jones, G. Ross, J. de Folter, M. Hartley, V. Uhlmann, A. Strange, Y. Schwab, E. Duke, C. Roufosse, and L. Collinson. 2025. A nanopathology pipeline for clinical research across scales using human tissue. medRxiv. 2025.08.29.25334660-2025.08.29.25334660. doi:10.1101/2025.08.29.25334660.

Bushong, E.A., D.D. Johnson, K.Y. Kim, M. Terada, M. Hatori, S.T. Peltier, S. Panda, A. Merkle, and M.H. Ellisman. 2015. X-ray microscopy as an approach to increasing accuracy and efficiency of serial block-face imaging for correlated light and electron microscopy of biological specimens. Microsc. Microanal. Off. J. Microsc. Soc. Am. Microbeam Anal. Soc. Microsc. Soc. Can. 21:231–238. doi:10.1017/S1431927614013579.

Darif, N., M. Rheinnecker, K. Hildenbrand, T. Chookajorn, L.P. Dorner, J.K. Hériché, S. Henriksson, C. Funaya, F. Hentzschel, L. Sandblad, O. Billker, Y. Schwab, and F. Frischknecht. 2026. Cellular Hallmarks From Volume Electron Microscopy Reveal Developmental Progression of Plasmodium Ookinetes. Adv. Sci. 13:e08250–e08250. doi:10.1002/ADVS.202508250.

Deerinck, T.J., E. A. Bushong, A. Thor, and M.H. Ellisman. 2010. NCMIR methods for 3D EM: A new protocol for preparation of biological specimens for serial block face scanning electron microscopy. 6–8.

Denk, W., and H. Horstmann. 2004. Serial Block-Face Scanning Electron Microscopy to Reconstruct Three-Dimensional Tissue Nanostructure. PLOS Biol. 2:e329–e329. doi:10.1371/JOURNAL.PBIO.0020329.

Fischler, M.A., and R.C. Bolles. 1981. Random sample consensus: A Paradigm for Model Fitting with Applications to Image Analysis and Automated Cartography. Commun. ACM. 24:381–395. doi:10.1145/358669.358692.

Handschuh, S., N. Baeumler, T. Schwaha, and B. Ruthensteiner. 2013. A correlative approach for combining microCT, light and transmission electron microscopy in a single 3D scenario. Front. Zool. 10:44. doi:10.1186/1742-9994-10-44.

Hoffman, D.P., G. Shtengel, C.S. Xu, K.R. Campbell, M. Freeman, L. Wang, D.E. Milkie, H.A. Pasolli, N. Iyer, J.A. Bogovic, D.R. Stabley, A. Shirinifard, S. Pang, D. Peale, K. Schaefer, W. Pomp, C.L. Chang, J. Lippincott-Schwartz, T. Kirchhausen, D.J. Solecki, E. Betzig, and H.F. Hess. 2020. Correlative three-dimensional super-resolution and block-face electron microscopy of whole vitreously frozen cells. Science. 367. doi:10.1126/SCIENCE.AAZ5357/SUPPL_FILE/AAZ5357S7.MOV.

Karreman, M.A., L. Mercier, N.L. Schieber, G. Solecki, G. Allio, F. Winkler, B. Ruthensteiner, J.G. Goetz, and Y. Schwab. 2016. Fast and precise targeting of single tumor cells in vivo by multimodal correlative microscopy. J. Cell Sci. 129:444–456. doi:10.1242/JCS.181842/-/DC1.

Kittelmann, M., C. Hawes, and L. Hughes. 2016. Serial block face scanning electron microscopy and the reconstruction of plant cell membrane systems. J. Microsc. 263:200–211. doi:10.1111/JMI.12424.

Klein, S., M. Staring, K. Murphy, M.A. Viergever, and J. Pluim. 2010. elastix: A Toolbox for Intensity-Based Medical Image Registration. IEEE Trans. Med. Imaging. 29:196–205. doi:10.1109/TMI.2009.2035616.

Kremer, A., E. van Hamme, J. Bonnardel, P. Borghgraef, C.J. Guérin, M. Guilliams, and S. Lippens. 2021. A workflow for 3D-CLEM investigating liver tissue. J. Microsc. 281:231–242. doi:10.1111/JMI.12967.

Kremer, J.R., D.N. Mastronarde, and J.R. McIntosh. 1996. Computer visualization of three-dimensional image data using IMOD. J. Struct. Biol. 116:71–76. doi:10.1006/JSBI.1996.0013.

Laundon, D., O.L. Katsamenis, J. Thompson, P. Goggin, D.S. Chatelet, P. Chavatte-Palmer, N.J. Gostling, and R.M. Lewis. 2023. Correlative multiscale microCT-SBF-SEM imaging of resin-embedded tissue. Methods Cell Biol. 177:241–267. doi:10.1016/BS.MCB.2023.01.014.

Lowe, D.G. 2004. Distinctive Image Features from Scale-Invariant Keypoints. Int. J. Comput. Vis. 60:91–110.

Maco, B., A. Holtmaat, M. Cantoni, A. Kreshuk, C.N. Straehle, F.A. Hamprecht, and G.W. Knott. 2013. Correlative in vivo 2 photon and focused ion beam scanning electron microscopy of cortical neurons. PloS One. 8:e57405. doi:10.1371/journal.pone.0057405.

Meechan, K., W. Guan, A. Riedinger, V. Stankova, A. Yoshimura, R. Pipitone, A. Milberger, H. Schaar, I. Romero-Brey, R. Templin, C.J. Peddie, N.L. Schieber, M.L. Jones, L. Collinson, and Y. Schwab. 2022. Crosshair, semi-automated targeting for electron microscopy with a motorised ultramicrotome. eLife. 11. doi:10.7554/ELIFE.80899.

Moore, J., D. Basurto-Lozada, S. Besson, J. Bogovic, J. Bragantini, E.M. Brown, J.M. Burel, X. Casas Moreno, G. de Medeiros, E.E. Diel, D. Gault, S.S. Ghosh, I. Gold, Y.O. Halchenko, M. Hartley, D. Horsfall, M.S. Keller, M. Kittisopikul, G. Kovacs, A. Küpcü Yoldaş, K. Kyoda, A. le Tournoulx de la Villegeorges, T. Li, P. Liberali, D. Lindner, M. Linkert, J. Lüthi, J. Maitin-Shepard, T. Manz, L. Marconato, M. McCormick, M. Lange, K. Mohamed, W. Moore, N. Norlin, W. Ouyang, B. Özdemir, G. Palla, C. Pape, L. Pelkmans, T. Pietzsch, S. Preibisch, M. Prete, N. Rzepka, S. Samee, N. Schaub, H. Sidky, A.C. Solak, D.R. Stirling, J. Striebel, C. Tischer, D. Toloudis, I. Virshup, P. Walczysko, A.M. Watson, E. Weisbart, F. Wong, K.A. Yamauchi, O. Bayraktar, B.A. Cimini, N. Gehlenborg, M. Haniffa, N. Hotaling, S. Onami, L.A. Royer, S. Saalfeld, O. Stegle, F.J. Theis, and J.R. Swedlow. 2023. OME-Zarr: a cloud-optimized bioimaging file format with international community support. Histochem. Cell Biol. 160:223–251. doi:10.1007/S00418-023-02209-1/FIGURES/15.

Peddie, C.J., and L.M. Collinson. 2014. Exploring the third dimension: Volume electron microscopy comes of age. Micron. 61:9–19. doi:10.1016/J.MICRON.2014.01.009.

Peddie, C.J., C. Genoud, A. Kreshuk, K. Meechan, K.D. Micheva, K. Narayan, C. Pape, R.G. Parton, N.L. Schieber, Y. Schwab, B. Titze, P. Verkade, A. Weigel, and L.M. Collinson. 2022. Volume electron microscopy. Nat. Rev. Methods Primer. 2:51. doi:10.1038/S43586-022-00131-9.

Randles, M.J., S. Collinson, T. Starborg, A. Mironov, M. Krendel, E. Königshausen, L. Sellin, I.S.D. Roberts, K.E. Kadler, J.H. Miner, and R. Lennon. 2016. Three-dimensional electron microscopy reveals the evolution of glomerular barrier injury. Sci. Rep. 6. doi:10.1038/SREP35068.

Ronchi, P., G. Mizzon, P. Machado, E. D’Imprima, B.T. Best, L. Cassella, S. Schnorrenberg, M.G. Montero, M. Jechlinger, A. Ephrussi, M. Leptin, J. Mahamid, and Y. Schwab. 2021. High-precision targeting workflow for volume electron microscopy. J. Cell Biol. 220:e202104069–e202104069. doi:10.1083/jcb.202104069.

Russell, M.R.G., T.R. Lerner, J.J. Burden, D.O. Nkwe, A. Pelchen-Matthews, M.C. Domart, J. Durgan, A. Weston, M.L. Jones, C.J. Peddie, R. Carzaniga, O. Florey, M. Marsh, M.G. Gutierrez, and L.M. Collinson. 2017. 3D correlative light and electron microscopy of cultured cells using serial blockface scanning electron microscopy. J. Cell Sci. 130:278–291. doi:10.1242/JCS.188433/VIDEO-5.

Schmidt, H., A. Gour, J. Straehle, K.M. Boergens, M. Brecht, and M. Helmstaedter. 2017. Axonal synapse sorting in medial entorhinal cortex. Nat. 2017 5497673. 549:469–475. doi:10.1038/nature24005.

Sengle, G., S.F. Tufa, L.Y. Sakai, M.A. Zulliger, and D.R. Keene. 2013. A correlative method for imaging identical regions of samples by micro-CT, light microscopy, and electron microscopy: Imaging adipose tissue in a model system. J. Histochem. Cytochem. 61:263–271. doi:10.1369/0022155412473757/SUPPL_FILE/10.1369_0022155412473757_SUPPLEMENTAL_MATERIAL.PDF.

Sofroniew, N., T. Lambert, G. Bokota, J. Nunez-Iglesias, P. Sobolewski, A. Sweet, L. Gaifas, K. Evans, A. Burt, D. Doncila Pop, K. Yamauchi, M. Weber Mendonça, J. Rodríguez-Guerra, L. Liu, G. Buckley, W.-M. Vierdag, A. Anderson, T. Monko, C. Willing, L. Royer, A. Can Solak, K.I.S. Harrington, J. Abramo, J. Ahlers, S. Ajina, D. Althviz Moré, O. Amsalem, E. Andò, A. Annex, C. Aronssohn, F. Balzaretti, P. Boone, K. Bestak, J. Bragantini, D. Bunten, M. Bussonnier, C. Caporal, M. Chazotte, I. Coccimiglio, Z. Čočková, J. Eglinger, A. Eisenbarth, J. Freeman, Y. Fukai T., C. Gohlke, K. Gunalan, Y.O. Halchenko, H. Har-Gil, M. Harfouche, V. Hilsenstein, K. Hutchings, H. Kawai, R. Kozar, J. Lauer, S. Le Meur-Diebolt, G. Lichtner, H. Liu, Z. Liu, A. Lowe, L. Marconato, S. Martin, A. McGovern, L. Migas, N. Miller, S. Miñano, H. Muñoz, J.-H. Müller, C. Nauroth-Kreß, H.A. Obenhaus, D. Palecek, C. Pape, E. Perlman, R.P. Theart, K. Pevey, G. Peña-Castellanos, A. Pierré, D. Pinto, C.M. Rodríguez-Reza, D. Ross, C.T. Russell, J. Ryan, G. Selzer, M. Smith, P. Smith, K. Sofiiuk, J. Soltwedel, D. Stansby, J. Vanaret, P. Wadhwa, M. Weigert, J. Windhager, P. Winston, Q. Yu, L. Zhang, R. Zhao, G. Witz, and M. Leomil Zoccoler. 2022. napari: a multi-dimensional image viewer for Python. Zenodo. doi:10.5281/ZENODO.19183461.

Svara, F., D. Förster, F. Kubo, M. Januszewski, M. Dal Maschio, P.J. Schubert, J. Kornfeld, A.A. Wanner, E. Laurell, W. Denk, and H. Baier. 2022. Automated synapse-level reconstruction of neural circuits in the larval zebrafish brain. Nat. Methods. 19:1357–1366. doi:10.1038/s41592-022-01621-0.

Titze, B., and C. Genoud. 2016. Volume scanning electron microscopy for imaging biological ultrastructure. Biol. Cell. 108:307–323. doi:10.1111/BOC.201600024.

Titze, B., C. Genoud, and R.W. Friedrich. 2018. SBEMimage: Versatile Acquisition Control Software for Serial Block-Face Electron Microscopy. Front. Neural Circuits. 12:383558–383558. doi:10.3389/FNCIR.2018.00054/BIBTEX.

Vergara, H.M., C. Pape, K.I. Meechan, V. Zinchenko, C. Genoud, A.A. Wanner, K.N. Mutemi, B. Titze, R.M. Templin, P.Y. Bertucci, O. Simakov, W. Dürichen, P. Machado, E.L. Savage, L. Schermelleh, Y. Schwab, R.W. Friedrich, A. Kreshuk, C. Tischer, and D. Arendt. 2021. Whole-body integration of gene expression and single-cell morphology. Cell. 184:4819–4837.e22. doi:10.1016/J.CELL.2021.07.017.

Wanner, A.A., C. Genoud, T. Masudi, L. Siksou, and R.W. Friedrich. 2016. Dense EM-based reconstruction of the interglomerular projectome in the zebrafish olfactory bulb. Nat. Neurosci. 2016 196. 19:816–825. doi:10.1038/nn.4290.

Zhang, Y., T. Ackels, A. Pacureanu, M.C. Zdora, A. Bonnin, A.T. Schaefer, and C. Bosch. 2022. Sample Preparation and Warping Accuracy for Correlative Multimodal Imaging in the Mouse Olfactory Bulb Using 2-Photon, Synchrotron X-Ray and Volume Electron Microscopy. Front. Cell Dev. Biol. 10:880696–880696. doi:10.3389/FCELL.2022.880696/TEXT.

Zheng, Z., J.S. Lauritzen, E. Perlman, C.G. Robinson, M. Nichols, D. Milkie, O. Torrens, J. Price, C.B. Fisher, N. Sharifi, S.A. Calle-Schuler, L. Kmecova, I.J. Ali, B. Karsh, E.T. Trautman, J.A. Bogovic, P. Hanslovsky, G.S.X.E. Jefferis, M. Kazhdan, K. Khairy, S. Saalfeld, R.D. Fetter, and D.D. Bock. 2018. A Complete Electron Microscopy Volume of the Brain of Adult Drosophila melanogaster. Cell. 174:730–743.e22. doi:10.1016/J.CELL.2018.06.019.

