## Supplemental figures for "iSBEM: An Open-Source Workflow for Automated ROI Targeting in Volume Electron Microscopy"

### Supplementary figures

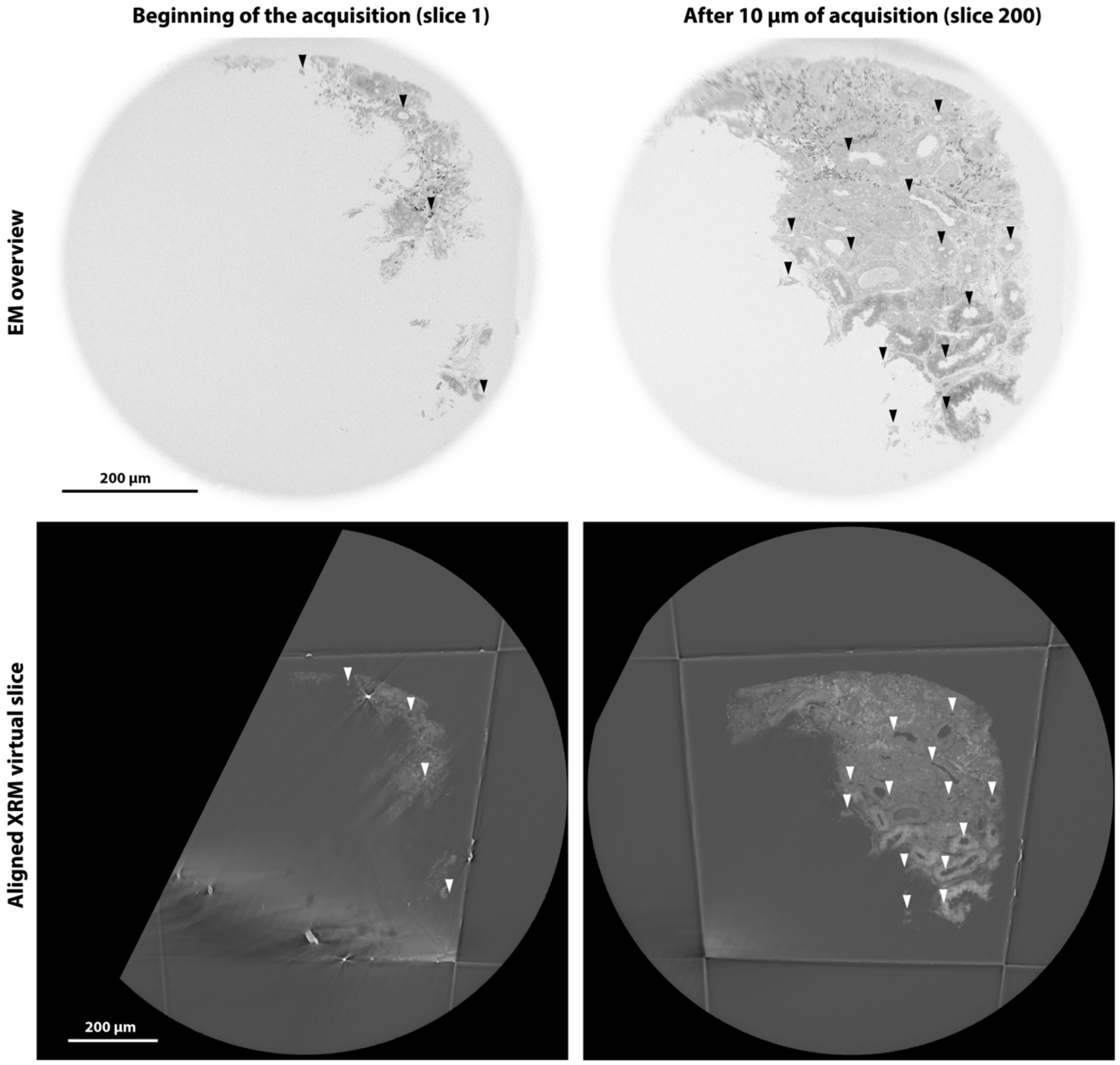


**Figure S1. Registration features**

The figure shows EM overviews (top panel) and corresponding XRM virtual slices after registration. Arrowheads highlight examples of anatomical features that can be used as landmarks for the registration. At the beginning of the acquisition, a small fraction of the tissue is exposed at the surface of the block and therefore only few features are available for the registration, often resulting in low accuracy. This becomes easier when more of the tissue is exposed.  Therefore, we routinely planned a registration checkpoint after 10 $\mu$m of acquisition.

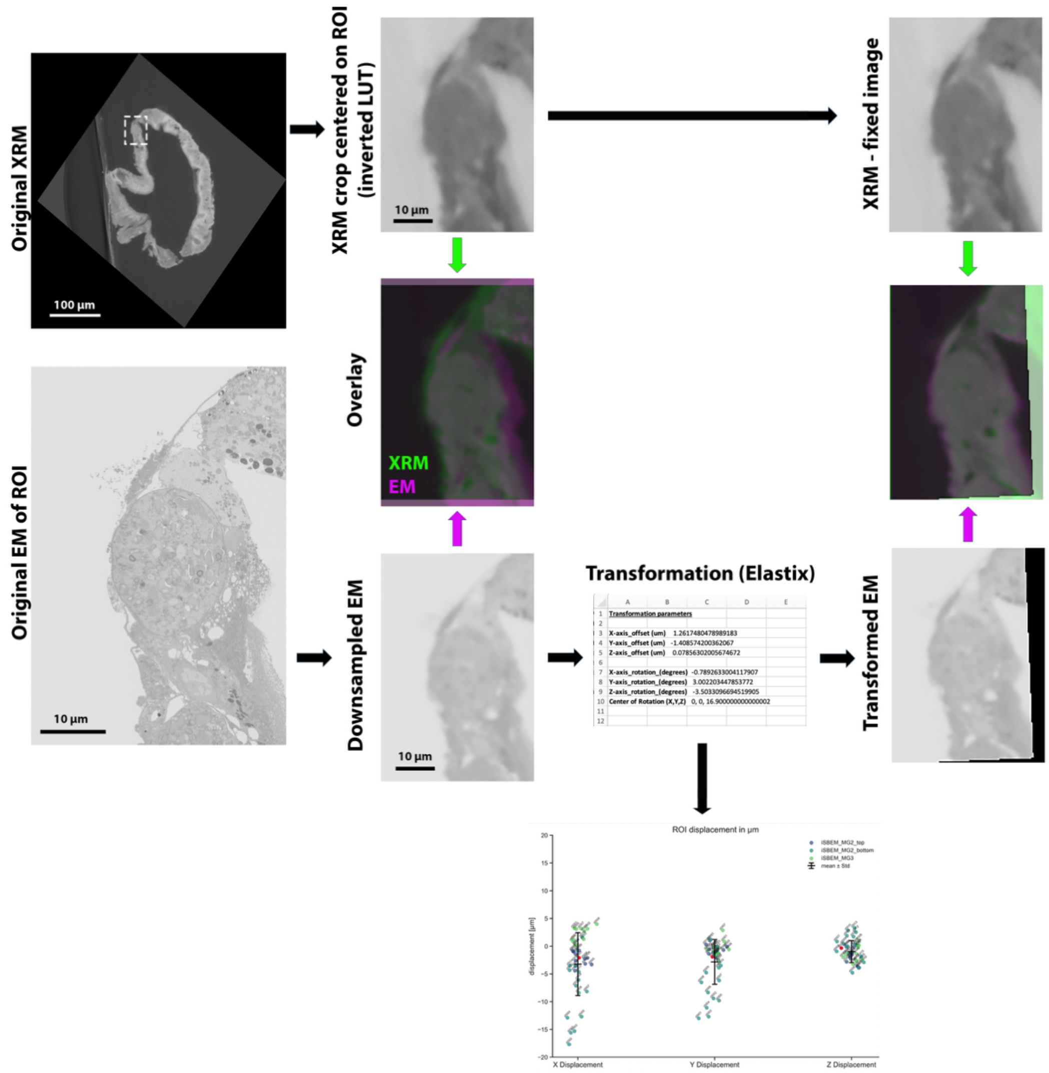


**Figure S2. Workflow of the strategy used to measure the registration accuracy (ROI displacement).**

Original XRM images are cropped around a segmented ROI with the same XYZ dimensions as the EM volume. Image contrast is inverted, to match the EM Look-Up Table (LUT). Original EM volumes are down-sampled to match the same resolution of the XRM (isotropic 0.65 µm). Overlay of the resulting volumes shows a displacement. The 2 volumes are then registered using Elastix allowing rigid transformation of one of the volumes (the down-sampled EM). Linear displacements as well as rotations in all axes are output by the software and are used to generate the displacement plots.

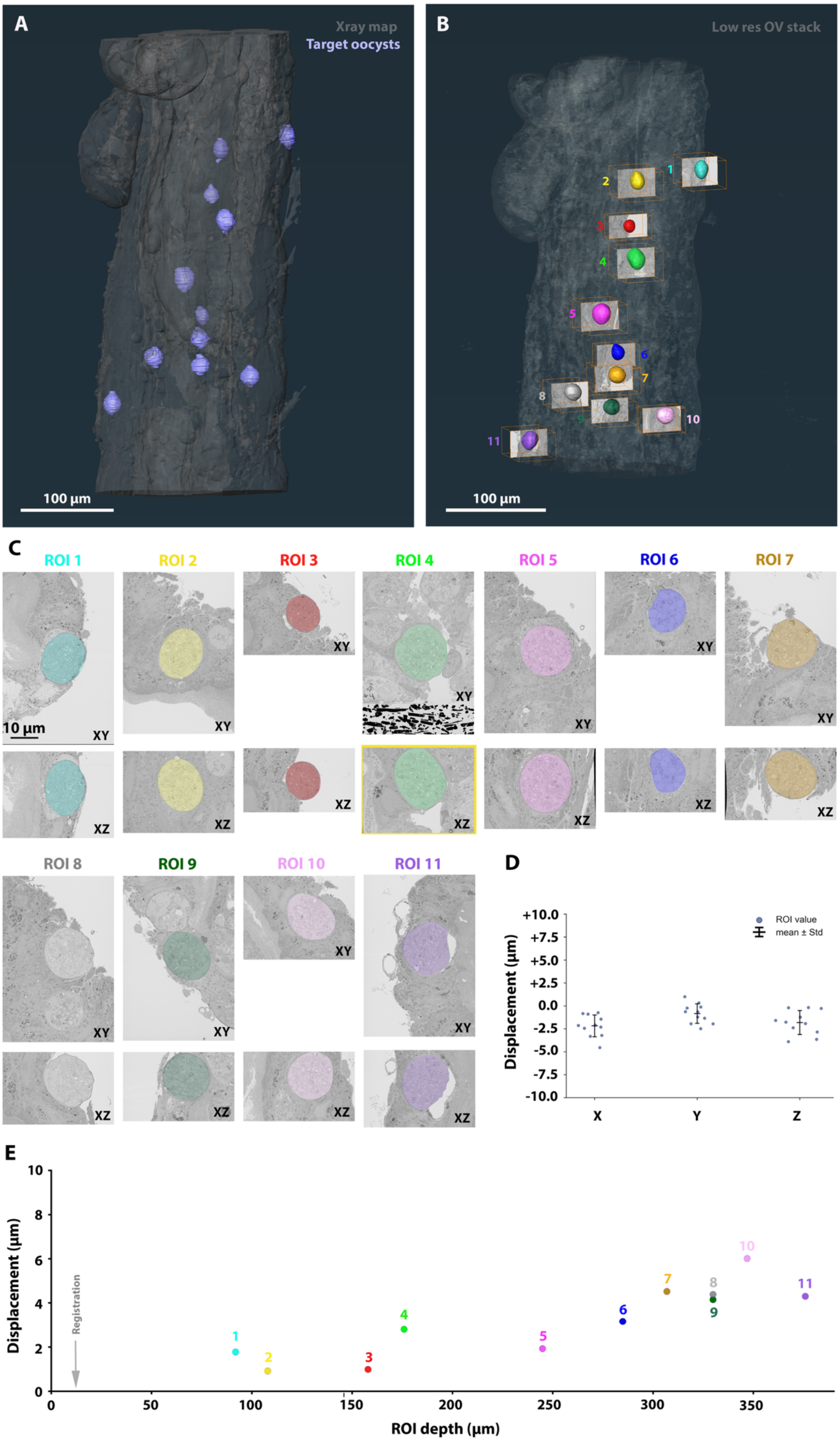

A
Xray map
Target oocysts
100 µm
B
Low res OV stack
1 2 3 4 5 6 7 8 9 10 11
100 µm
C
ROI 1
ROI 2
ROI 3
ROI 4
ROI 5
ROI 6
ROI 7
ROI 8
ROI 9
ROI 10
ROI 11
XY
XZ
10 µm
D
Displacement (µm)
ROI value
mean ± Std
X
Y
Z
E
Displacement (µm)
Registration
ROI depth (µm)

**Figure S3. Fully automated targeted acquisition of malaria oocysts in an infected mosquito midgut using iSBEM. Example 1.**

A. Volume rendering of the mosquito midgut (grey) from the XRM volume with the segmented oocyst targets (lavender). To add a margin of error in Z for the acquisition, the segmentation was extended in Z, giving the round oocysts an elongated shape.

B. Volume rendering of the mosquito midgut generated from SBF-SEM OVs (in grey transparent), overlaid with an XZ plane from each of the acquired high resolution EM volumes for each ROI and a 3D rendering of the oocysts.

C. Central cross section of all acquired ROI volumes in XY and XZ, with the targeted oocyst overlaid in colors. The central position of the oocysts in both axes shows the high precision of the targeting. A yellow frame highlights the target that was not entirely acquired. In this case, the oocyst in ROI4 was partially missed at the top, resulting in the loss of a very small fraction of volume.

D. Displacement (Euclidean distance) obtained for all ROIs in individual axes. Each dot represents an ROI.

E. Targeting accuracy measurement. The EM volume of each ROI was registered to the corresponding XRM crop around the segmented target (see Figure S2). The transformation vector is displayed in the chart (ROI displacement) as a function of the ROI depth. The arrow shows the registration point.

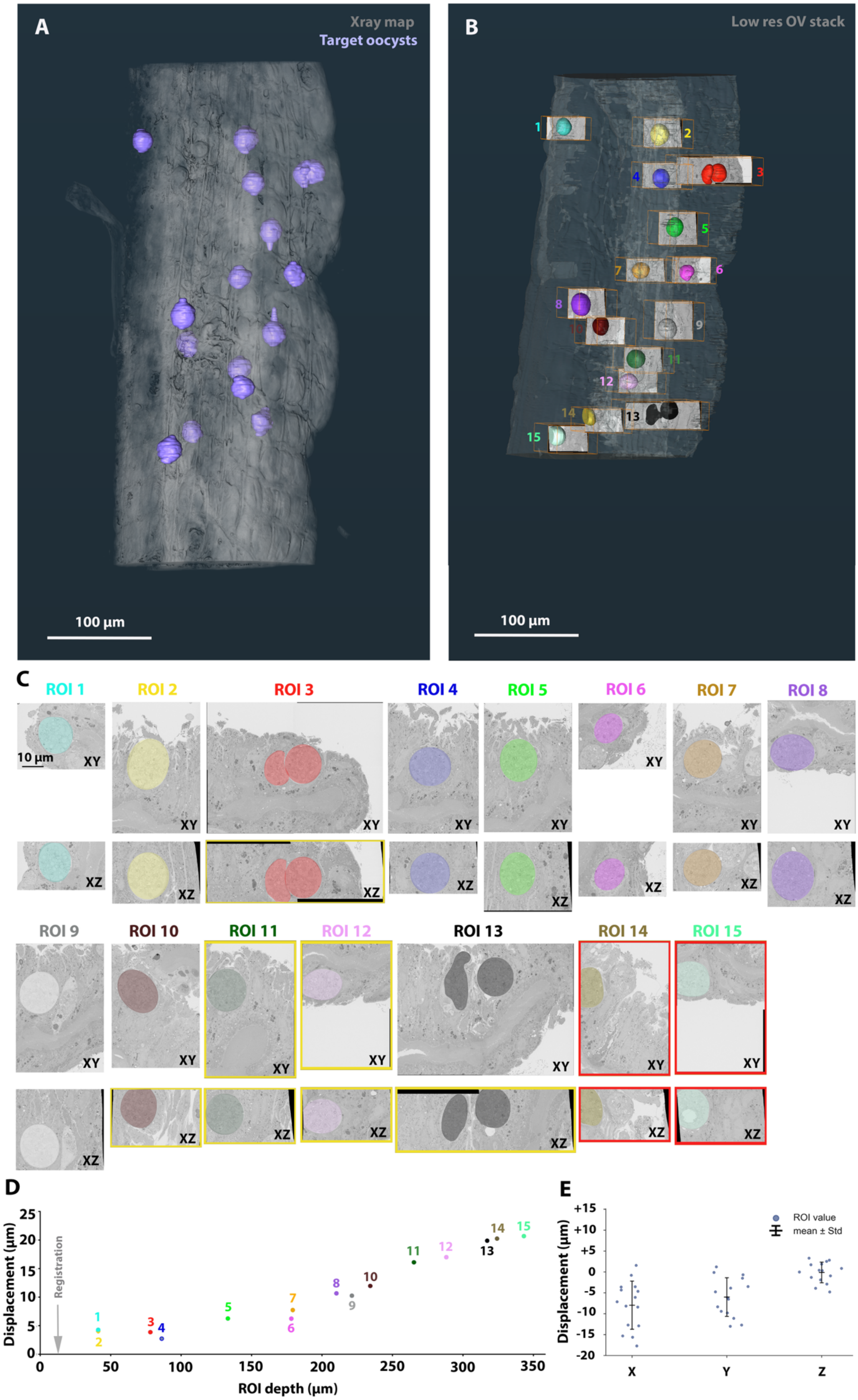
A
Xray map
Target oocysts
100 µm
B
Low res OV stack
100 µm
C
ROI 1
ROI 2
ROI 3
ROI 4
ROI 5
ROI 6
ROI 7
ROI 8
10 µm
XY
XZ
ROI 9
ROI 10
ROI 11
ROI 12
ROI 13
ROI 14
ROI 15
D
Displacement (µm)
Registration
ROI depth (µm)
E
ROI value
mean ± Std
Displacement (µm)
X
Y
Z

**Figure S4. Fully automated targeted acquisition of malaria oocysts in an infected mosquito midgut using iSBEM. Example 2.**

A. Volume rendering of the mosquito midgut (grey) from the XRM volume with the segmented oocyst targets (lavender). To add a margin of error in Z for the acquisition, the segmentation was extended in Z, giving the round oocysts a spheroid shape with elongated ends.

B. Volume rendering of the mosquito midgut generated from SBF-SEM OVs (in grey transparent), with overlays of the acquired high resolution volumes for each ROI. Shown is a XZ plane of the acquired high resolution EM volumes of each ROI and a rendering of the targeted oocysts.

C. Central cross section of all acquired ROI volumes in XY and XZ, with the targeted oocyst overlaid in colors. A yellow or red frame highlights the targets that were not entirely acquired. Yellow, minor volume loss; red, major volume loss.

D. Targeting accuracy measurement. The EM volume of each ROI was registered to the corresponding XRM crop around the segmented target (see Fig S2). The transformation vector is displayed in the plot (ROI displacement) as a function of the ROI depth. The arrow shows the registration point.

E. Displacement (Euclidean distance) obtained for all ROIs in individual axes. Each dot represents an ROI.

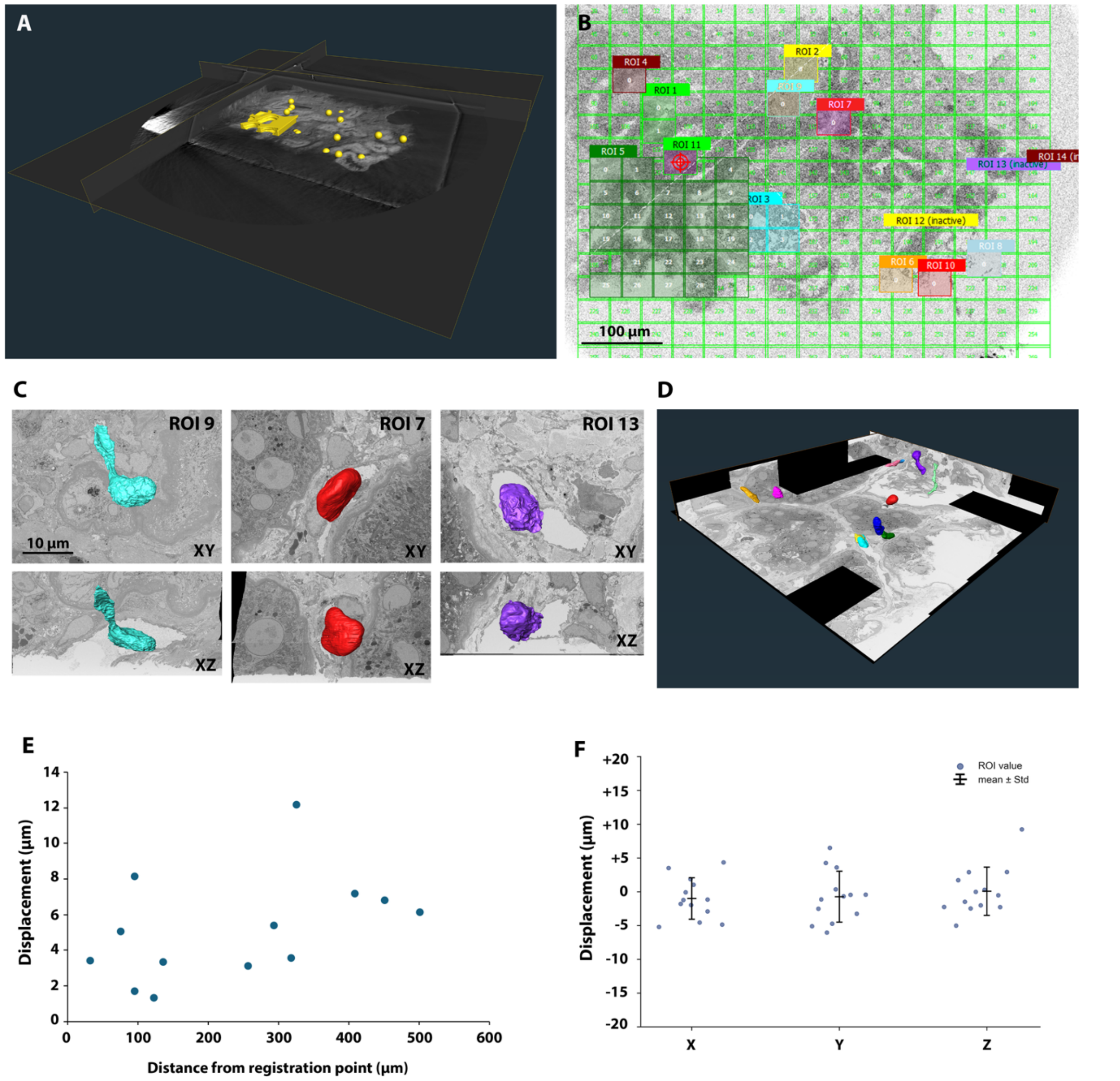


**<u>Figure S5. Fully automated targeting of immune cells in a human kidney biopsy. Example 2</u>**

A. 3D visualization of the XRM dataset overlaid with a mask of the ROIs to be targeted by SBF-SEM in yellow.

B. Snapshot from the SBEMimage interface, showing all the grids that were generated from the segmented ROIs. ROIs that are active at this point of the acquisition (1,2,3,4,5,6,7,8,9,10,11) are highlighted.

C. Examples of three intracapillary immune cells targeted with the iSBEM workflow. Segmentation projected on an XY plane is shown in the top panels. Segmentation projected on the XZ plane in the lower ones.

D. 3D visualization of the EM volume of a large biopsy region (ROI 5, including part of a glomerulus and some peritubular capillaries), after tile stitching and slice alignment. Immune cells in glomerular and peritubular capillaries are segmented and rendered in different colors.

E. Displacement in 3D as a function of the distance from the initial registration point (see materials and methods). Each dot represents a single cell. Compared to figure 4L, the displacements are smaller and the dependence of the offset on the distance from the registration point is less obvious, suggesting that the initial registration was more accurate in this case.

F. Displacement (Euclidean distance) obtained for all ROIs in individual axes. Each dot represents an ROI.

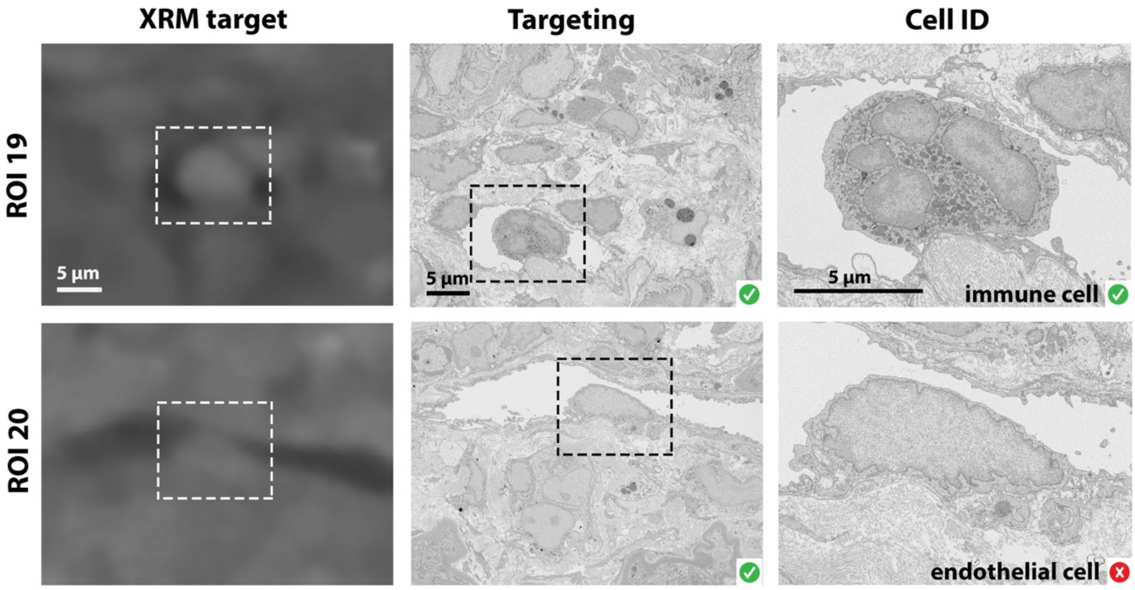


**Figure S6. Examples of targeting and immune cell identification accuracy of the workflow.**

Cells were identified as potential immune cells in the XRM volume (left panel, dashed boxes) and targeted for SBF-SEM using the iSBEM workflow. Targeting the cells of interest was successful, as shown in the central panel (targets in the black dashed boxes), but only the top one proved to be an immune cell, while the lower one was in fact an endothelial cell.
